# Nonlinear effects of intrinsic dynamics on temporal encoding in a model of avian auditory cortex

**DOI:** 10.1101/2020.07.28.225680

**Authors:** Christof Fehrman, Tyler D Robbins, C Daniel Meliza

## Abstract

Neurons exhibit diverse intrinsic dynamics, which govern how they integrate synaptic inputs to produce spikes. Intrinsic dynamics are often plastic during development and learning, but the effects of these changes on stimulus encoding properties are not well known. To examine this relationship, we simulated auditory responses to zebra finch song using a linear-dynamical cascade model, which combines a linear spectrotemporal receptive field with a dynamical, conductance-based neuron model, then used generalized linear models to estimate encoding properties from the resulting spike trains. We focused on the effects of a low-threshold potassium current (K_LT_) that is present in a subset of cells in the zebra finch caudal mesopallium and is affected by early auditory experience. We found that K_LT_ affects both spike adaptation and the temporal filtering properties of the receptive field. The direction of the effects depended on the temporal modulation tuning of the linear (input) stage of the cascade model, indicating a strongly nonlinear relationship. These results suggest that small changes in intrinsic dynamics in tandem with differences in synaptic connectivity can have dramatic effects on the tuning of auditory neurons.

**Author summary:** Experience-dependent developmental plasticity involves changes not only to synaptic connections, but to voltage-gated currents as well. Using biophysical models, it is straightforward to predict the effects of this intrinsic plasticity on the firing patterns of individual neurons, but it remains difficult to understand the consequences for sensory coding. We investigated this in the context of the zebra finch auditory cortex, where early exposure to a complex acoustic environment causes increased expression of a low-threshold potassium current. We simulated responses to song using a detailed biophysical model and then characterized encoding properties using generalized linear models. This analysis revealed that this potassium current has strong, nonlinear effects on how the model encodes the song’s temporal structure, and that the sign of these effects depend on the temporal tuning of the synaptic inputs. This nonlinearity gives intrinsic plasticity broad scope as a mechanism for developmental learning in the auditory system.

## Introduction

Neurons have diverse, nonlinear dynamics. Many brain regions contain multiple kinds of neurons with different spike waveforms and spiking patterns (Bal and Oertel, 2001; Sivaramakrishnan and Oliver, 2001; Ascoli et al., 2008), and there is substantial variation even within well-defined cell types (Toledo-Rodriguez et al., 2004; Schulz et al., 2006; Bomkamp et al., 2019). Intrinsic dynamics can be modified by activity and experience (Ross et al., 2017; Daou and Margoliash, 2020; Chen and Meliza, 2020), which may be an important mechanism for learning (Titley et al., 2017). This physiological diversity has been known for many decades (Llinás, 1988) and can be modeled on a detailed, biophysically realistic level (Padmanabhan and Urban, 2010; Tripathy et al., 2015), but our understanding of how intrinsic dynamics affect neural computations in many systems has remained surprisingly qualitative.

The complexity and nonlinearity of biophysical models makes it difficult to use them to explain higher-order processes in the brain, at what Marr (1982) termed the algorithmic and computational levels. A simple, single-compartmental model that can produce common physiological behaviors like bursting, adaptation, or rebound spiking, is a system of around ten or more nonlinear differential equations, with fifty or more parameters (e.g., Rothman and Manis, 2003; Meliza et al., 2014). These parameters correspond to specific aspects of the cell biology (such as membrane capacitance or sodium channel density), which makes them easy to interpret and, in some cases, possible to measure directly. However, the relationships between the parameters and the observable behaviors of the neuron are highly nonlinear, making it difficult to constrain them statistically. It is difficult and time-consuming to fit dynamical models to biological data (Druckmann et al., 2008; Van Geit et al., 2008; Toth et al., 2011; Vavoulis et al., 2012), and there is little consensus on the appropriate methods or even whether there are globally optimal solutions (Prinz et al., 2004). Moreover, access to the intracellular voltage is needed, through a sharp or patch electrode or using an optical sensor (Huys and Paninski, 2009), which greatly limits the number of neurons that can be modeled within the context of a circuit, and almost always requires the use of *ex vivo* preparations that cannot be presented with realistic stimuli.

As a consequence, many studies of function in neural systems have emphasized phenomenological models that omit most of the biophysical and dynamical features of spike generation in exchange for computational tractability (Keat et al., 2001; Jolivet et al., 2004; Kobayashi et al., 2009; Izhikevich, 2003). One of the simplest examples is the generalized linear model (GLM), which represents spiking as an inhomogeneous Poisson process with a conditional intensity that depends only on a linear function of the stimulus and spiking response in the recent past (Pillow et al., 2005). In contrast to more realistic models, the GLM is a staple of statistics, with a well-defined likelihood function that is concave everywhere, guaranteeing that a global optimum can be found (Paninski et al., 2004). The GLM also has established techniques for regularization, which is necessary when stimuli have naturalistic (i.e., highly correlated) distributions (Theunissen et al., 2001; Schwartz et al., 2006).

Because of its simplicity and probabilistic formulation, a GLM can be thought of as a representation of a neuron’s encoding properties; that is, an abstract view of how the cell transforms sensory stimuli into spike trains. Surprisingly, although GLMs have been successfully used to model encoding in a number of different sensory systems (Pillow et al., 2005; Calabrese et al., 2011), and there have been several studies using GLMs to predict and characterize more complex spiking models (Ostojic and Brunel, 2011; Pozzorini et al., 2015; Weber and Pillow, 2017), to our knowledge there has not been any attempt to relate the GLM to more detailed, dynamical models with realistic sensory inputs. As a result, it is difficult to predict how natural, pathological, or experience-dependent variations in voltage-gated channels are likely to affect sensory processing.

In this study, we examined the relationship between intrinsic dynamics and encoding properties in the context of auditory processing in songbirds. Encoding models, including GLMs, have been employed extensively to study this system (Theunissen et al., 2000; Sen et al., 2001; Nagel and Doupe, 2008; Woolley et al., 2009; Calabrese et al., 2011), but until recently, there have been no data on the intracellular physiology of the constituent neurons. Using whole-cell patch recordings from slices, we have found that the caudal mesopallium (CM), a cortical-level auditory area (Wang et al., 2010; Jarvis et al., 2013), has diverse, experience-dependent intrinsic dynamics (Chen and Meliza, 2018, 2020). Most of the putatively excitatory neurons fire repetitively when depolarized, but a substantial fraction only fire at stimulus onsets. This phasic firing behavior is correlated with strong outward rectification that activates at low voltages, and it can be pharmacologically converted to tonic firing by blocking low-threshold potassium currents (K_LT_). The proportion of phasic neurons changes over development, reaching a peak around the age zebra finches begin to memorize songs, but only in birds exposed to a complex acoustic environment. This experience-dependent plasticity is correlated with changes in the expression of Kv1.1, a low-threshold potassium channel (Chen and Meliza, 2020).

The dependence of phasic firing on auditory experience suggests that intrinsic plasticity (i.e., a change in the expression or properties of voltage-gated currents, rather than synaptic currents) plays a critical role in development for songbirds, but for all the reasons noted above, the functional significance remains unclear. Here, we took a simulation-based approach to ask how changing the magnitude of low-threshold potassium currents in a dynamical model would affect encoding properties, as estimated with a GLM. We simulated auditory responses using a linear-dynamical cascade model (Bjoring and Meliza, 2019), which combines a linear spectrotemporal receptive field (RF) with a single-compartment biophysical model (Fig. 1A). The linear stage of the model consists of representative RFs based on the data and parametric model of Woolley et al. (2009), which are convolved with spectrograms of zebra finch song to generate an external driving current. Conceptually, this current represents a linear approximation of the summation and filtering performed by the neuron’s dendrites on excitatory and inhibitory synaptic inputs. The biophysical model we used includes sodium, high-threshold potassium, transient (A-type) potassium, low-threshold potassium, and hyperpolarization-activated (h-type) currents, and it can reproduce the responses of phasic and tonic CM neurons to step and broadband current stimuli (Chen and Meliza, 2018). As shown previously, phasic firing in this model depends on a single parameter that governs the maximal conductance of the low-threshold potassium current (*g_KLT_*) (Fig. 1B–C). We used the spike trains produced by these simulations to fit GLMs (Fig. 2) and then compared estimates for the RF and spike-history parameters to determine how K_LT_ influenced how the model was encoding the acoustic structure of the stimulus.

**Fig. 1.**
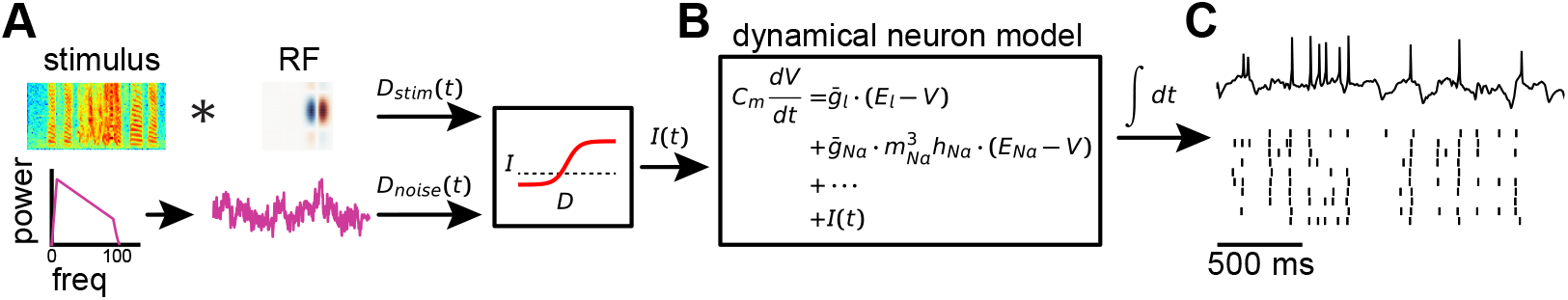
Linear-dynamical cascade model. (A) The linear stage of the model consists of the convolution of a stimulus with a receptive field. The output of the convolution (*D*_stim_(*t*)) is combined with a stimulus-independent noise signal (*D*_noise_(*t*)) with a 1/ *f* spectral distribution. The sum of *D*_noise_(*t*) and *D*_stim_(*t*) is converted to the input current *I*(*t*) using a static nonlinearity, ensuring that the model voltage remains within biologically realistic bounds. (B) *I*(*t*) enters into the biophysical stage, which models membrane voltage dynamics as a system of ordinary differential equations. (C) The model is numerically integrated to produce a simulated voltage trace. Multiple trials are simulated by keeping *D_stim_*(*t*) the same from trial to trial, while drawing new values for *D*_noise_(*t*).

**Fig. 2.**
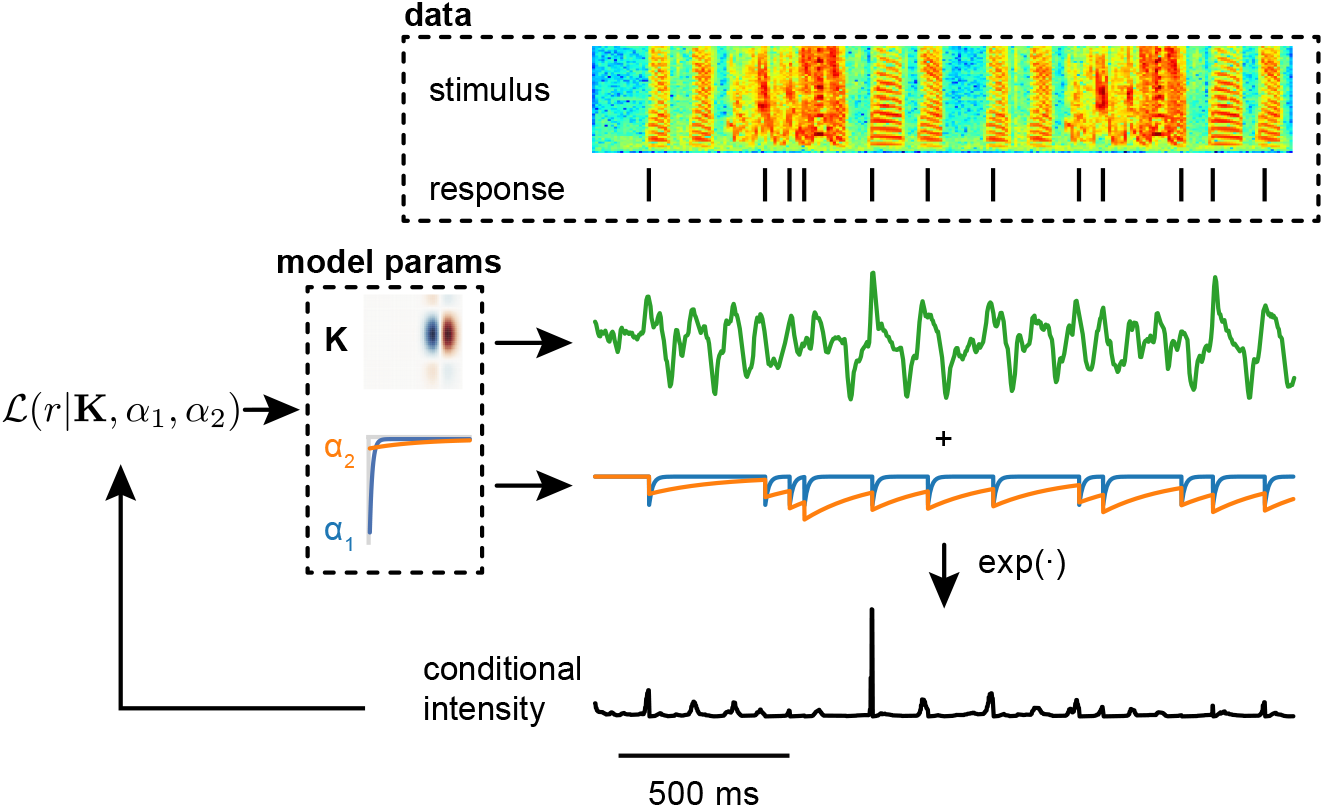
Schematic of parameter estimation for generalized linear model. The data to be fit comprise a stimulus, which can be a univariate time series or a multivariate spectrogram (as shown here), and a spiking response. The model represents the response as an inhomogeneous Poisson process with a conditional intensity that depends on the convolution of the stimulus with a receptive field (**K**) and the convolution of the response with a spike-history filter, which was parameterized as the sum of two exponential decays representing short-term (*α*_1_) and long-term (*α*_2_) adaptation or facilitation. Not shown is a constant offset *ω*, which governs the baseline probability of firing, such that higher values suppress the probability of spiking. These model parameters are estimated by regularized maximum likelihood.

## Results

### Univariate white-noise stimulus

As a proof of principle, we began with an example using a white-noise stimulus drawn from a univariate Gaussian distribution. The absence of temporal correlations in this stimulus is ideal for obtaining unbiased estimates of the GLM parameters, allowing us to determine how intrinsic dynamics affect encoding in a best-case scenario.

We generated data for fitting the GLM by providing 100 s of white noise as input to two linear-dynamical cascade (LDC) models that had the same RF but different dynamics. The dynamical stage of the model was based on our previous work in the zebra finch caudal mesopallium (Chen and Meliza, 2018; Bjoring and Meliza, 2019). The tonic model lacks K_LT_ and has a higher capacitance, whereas the phasic model includes K_LT_ and has a lower capacitance (see Methods for parameter values). These models reproduce the responses to step currents (Fig. 3A) and broadband currents seen in slices. Both LDC models produced similar responses to the white noise stimulus, but the phasic model tended to have narrower peaks of activity (Fig. 3B–C).

**Fig. 3.**
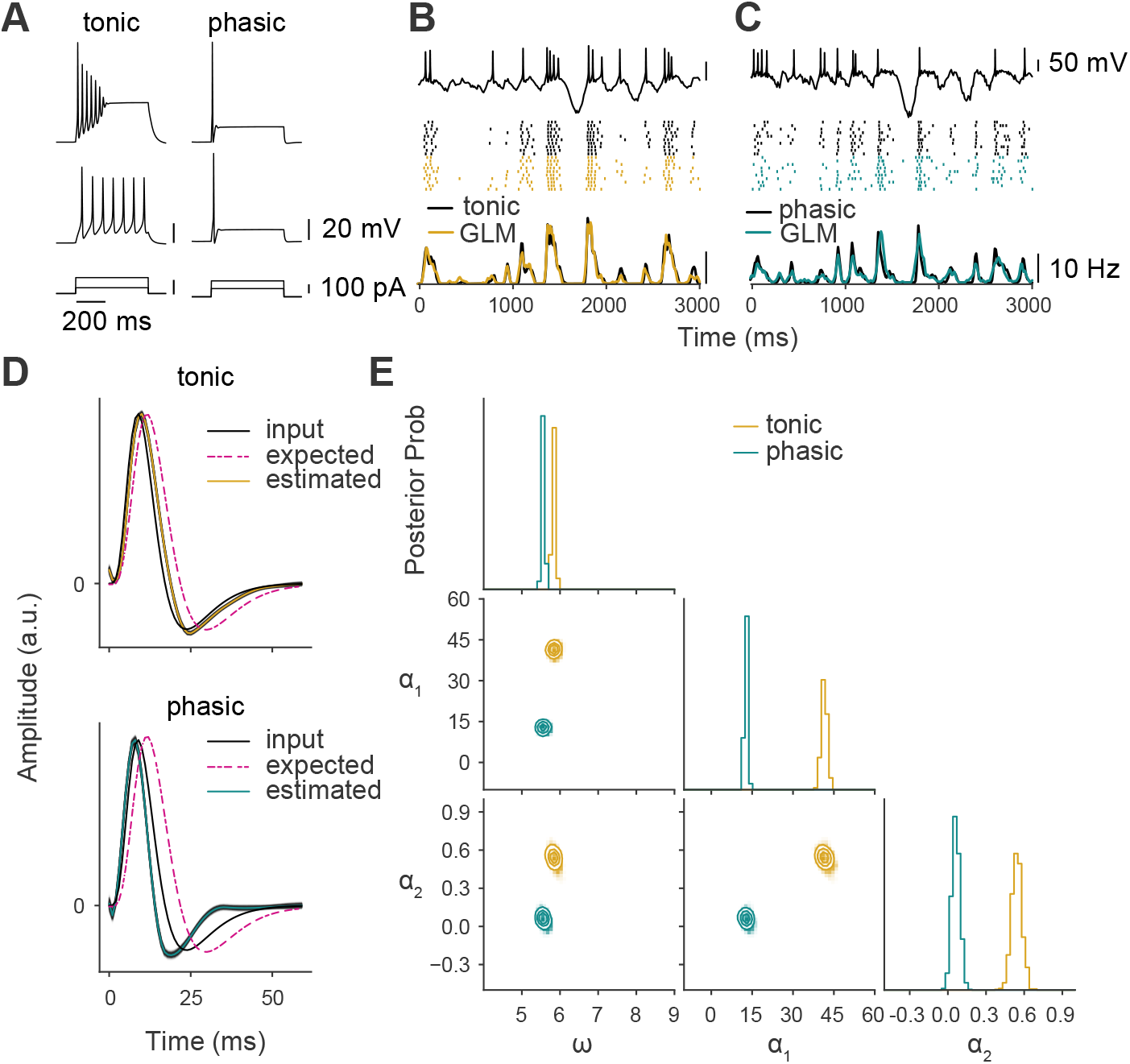
GLM estimates for exemplar tonic and phasic models with univariate white-noise stimulus. (A) Voltage responses of tonic and phasic models to high- and low-amplitude injected current steps (shown in bottom row). The tonic model exhibits depolarization block to strong currents but fires repetitively to weak currents, whereas the phasic model only fires a single spike to all suprathreshold current levels. (B) Top, response of the tonic dynamical model to a white-noise stimulus. The input RF is shown in D. Middle, raster plots of spike times from 10 trials with the same stimulus but varying *I*_noise_(*t*). Black ticks correspond to the output of the dynamical model and colored ticks are the predictions of a GLM fit to a different set of data from this model. Bottom, spike rate histograms (bin size = 10 ms) for 50 trials from the dynamical model (black) and the GLM (yellow). Only a subset of the full test data is shown. (C) Like B, but for the model with phasic dynamics. The stimulus, RF, and noise level were the same. (D) Estimated RFs from the GLMs compared to the input RF of the dynamical model. To indicate posterior uncertainty in the estimates, individual samples from the MCMC sampler are shown in light gray, and the median is overlaid in color. (E) Posterior distributions of baseline firing rate (*ω*) and spike-history filter parameters (*α*_1_ and *α*_2_). The top panels in each column show marginal distributions for individual parameters, and the panels in the lower left corner show joint distributions for each pair of parameters. Note that more positive values of *α*_1_ and *α*_2_ correspond to stronger adaptation (i.e., a negative correlation with past spiking).

In general, parameter estimates are only interpretable to the extent that the model is a good fit to the data. We checked the goodness of fit by comparing the responses of the LDC model and the fitted GLM to a new white-noise stimulus. The output of the GLM was an excellent prediction of the dynamical model’s response (Fig. 3B–C). Indeed, the correlations between the average firing rates for LDC data and GLM prediction (tonic: *r* = 0.96; phasic: *r* = 0.84) were comparable to the correlations between average rates of even and odd trials in the data (tonic: *r* = 0.94; phasic: *r* = 0.90)—as good as could be expected given the intrinsic variability of the data. Thus, at least for white-noise stimuli, the linear spike-history filter and static nonlinearity of the GLM can closely approximate the dynamical nonlinearity of a single-compartment biophysical model. This allows us to interpret the GLM parameters as meaningful descriptions of the encoding properties of the more complex model.

The LDC and GLM both have receptive fields that are convolved with the stimulus to produce a signal that modulates the probability of spiking. When a GLM is fit using data from an LDC model, we expect the estimated RF to resemble the RF used to generate the data, but not exactly.

Indeed, differences between the input and estimated RFs will reflect the effects of the intrinsic dynamics. One expected effect is from the filtering properties of the membrane. In the GLM, firing probability depends on a static, exponential function of the convolved stimulus (Fig. 2). In the LDC model, the output of the convolution enters as a current that contributes linearly to the derivative of the membrane voltage. The capacitance and conductance of the membrane act as an additional, lowpass filter, so we would expect the estimated RF to be a lowpass-filtered version of the input RF. In the time domain, the effect of the membrane would be to stretch the RF out in time. In fact, what we observed was that the estimated RFs were either very close to the input RF (Fig. 3D, top) or compressed in time (Fig. 3D, bottom), corresponding to a relative boosting of higher frequencies. This would not be possible for a model with a purely passive membrane; therefore, it must be the active, voltage-gated currents that are shifting the model’s temporal encoding properties. This temporal distortion, which is consistent with the bandpass characteristics of K_LT_ (Meng et al., 2012; Chen and Meliza, 2018), will be explored further in subsequent analyses.

Intrinsic dynamics also affected the spike-history filter. Unlike the RF, the parameters for the spike-history filter do not correspond to specific parameters in the LDC model; however, we expect them to reflect the effects of currents that are activated by spiking. As seen in Fig. 3E, the spike-history filter was stronger on both short (*α*_1_) and long (*α*_2_) timescales for data from the tonic model compared to the phasic one. The posterior uncertainty in these parameter estimates was low compared to the difference between dynamical models. This means that the spiking patterns produced by phasic and tonic cells are sufficiently different, at least for this kind of stimulus and amount of data, to observe changes in a single biophysical parameter.

### Multivariate birdsong stimulus

Having demonstrated that the GLM can be used to analyze the encoding properties of a dynamical model, we turned to a more realistic scenario using natural birdsong as the stimulus. The dynamics remained the same as in the white-noise case, but the linear stage was replaced with a spectrotemporal RF. The stimulus, which consisted of 40 s of song from multiple zebra finches, was converted to a spectrogram and convolved with the RF, summing across spectral channels. This produced a univariate time series that entered into the dynamics as an external current.

We used RFs that were representative of the diversity found in cortical-level auditory neurons. RF structure can be analyzed in terms of the modulation transfer function (MTF), a 2-D Fourier transform of the RF that shows its joint spectral and temporal tuning (Woolley et al., 2005). Most of the neurons in the zebra finch primary auditory pallium have MTFs with power along either the spectral or temporal axis, indicating that they can be tuned to narrow spectral bands or to rapid modulations of the temporal envelope, but only rarely to both (Woolley et al., 2009). This distribution is similar to the modulation spectrum of zebra finch song (Woolley et al., 2005) and at least partly reflects the statistics of early auditory experience (Moore and Woolley, 2019). Here, we simulated responses using 60 synthetic RFs drawn from this distribution (Woolley et al., 2009). Each RF was combined with the tonic and phasic dynamical models, so that we could quantify the effects of K_LT_ across RF types and determine if there was any interaction with RF structure.

As before, the simulated responses were used to estimate GLM parameters, but with two modifications that were necessitated by the statistics of the birdsong stimuli. Like many other natural stimuli, the amplitude envelope of birdsong is dominated by low frequencies (Singh and Theunissen, 2003). For our cascade model, these low-frequency temporal modulations result in long intervals when *I*(*t*) is strongly positive or negative, which in turn tends to drive the model to unrealistic voltage levels far outside the range that would be expected from the reversal potentials of typical synaptic channels. To address this issue, we introduced a compressive static nonlinearity that constrained the output of the convolution to biologically feasible values (see Methods). The second issue with stimuli dominated by low frequencies is a statistical one. As has been known for some time (Theunissen et al., 2000, 2001), estimating the parameters of receptive field models when the stimulus is highly autocorrelated can lead to numerical instability and overfitting. To address this issue, we used elastic-net regularization when estimating GLM parameters (see Methods).

We begin by examining three examples representative of the distribution. As will be seen, the temporal characteristics of the input RF have a consistent effect on encoding properties, so we have denoted these three examples in terms of their temporal modulation transfer functions (tMTFs): wideband (WB), bandpass-low (BP-L), and bandpass-high (BP-H). These categories reflect two parameters in the equation we used to generate RFs (see Methods). Wideband RFs have a temporal phase (*P_t_*) of zero, which results in only a single excitatory lobe in the temporal profile and broad tuning in the temporal modulation frequency domain. Bandpass RFs have a temporal phase of 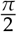, resulting in a suppressive/inhibitory lobe. BP-L and BP-H are distinguished by the frequency modulation parameter (Ω_*t*_), with lower values corresponding to a broader temporal profile and tuning to slower modulations. As seen in Fig. 4A–F, the fitted GLMs had good predictive performance for both the phasic and tonic models and across all three input RFs, with high correlations between the spike rate histograms produced by the LDC and GL models to a novel birdsong stimulus. Thus, even with many more parameters and an autocorrelated stimulus, the GLM is still a good tool for analyzing the encoding properties of the dynamical models.

**Fig. 4.**
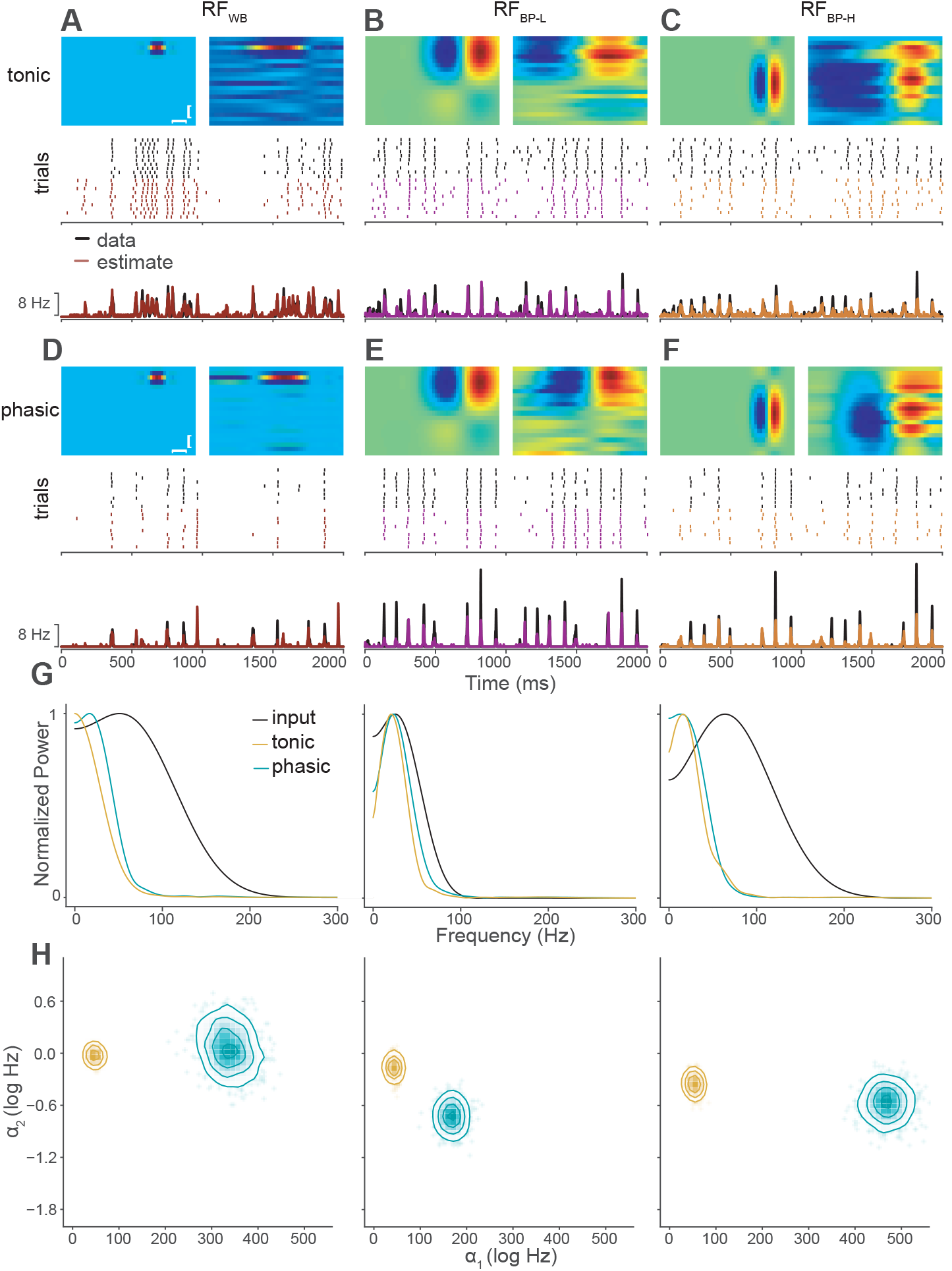
GLM estimates for exemplar tonic and phasic models with zebra finch song stimuli. (A) Receptive field parameters and responses for a model with tonic dynamics and a spectrally narrowband, temporally wideband RF. Top left, input RF in the LDC model. Top right, estimated RF from GLM. The vertical scale bar denotes 1 kHz and the horizontal 5 ms. Note the temporal smearing and the broad suppression at longer lags in the estimated RF. Middle, examples of spiking responses to zebra finch song from the LDC model (top, black ticks) and the fitted GLM (bottom, red). Bottom, corresponding spike rate histograms (50 trials) for the LDC and GLM (product-moment correlation: r_WB_ = 0.87). (B–C) RFs and responses for models with tonic dynamics and BP-L (B) or BP-H RFs (C), same format as in (A). The GLM accurately predicted the firing rate of the LDC for these parameter values (r_BP-L_ = 0.94, r_BP-H_ = 0.86). (D–F) RFs and responses for models with the same RFs as in (A–C), but with phasic dynamics (r_WB_ = 0.78, r_BP-L_ = 0.90, r_BP-H_ = 0.85). All prediction correlations were high considering the underlying spiking variability in the even and odd trials of the LDC (product-moment correlations: tonic_WB_ = 0.92, tonic_BP-L_ = 0.85, tonic_BP-H_ = 0.82; phasic_WB_ = 0.91, phasic_BP-L_ = 0.93, phasic_BP-H_ = 0.91). More detailed plots for each of the six example models can be found in S1 Text. (G) Temporal MTFs of input RFs, tonic model estimates, and phasic model estimates for each of the three input RFs. Power is normalized relative to the peak for each spectrum. The change in power at low frequencies, quantified as Δ*l* (see Methods) was –0.08, 0.44, and –0.15 for tonic models and –0.03, 0.30, and –0.33 for phasic models. (H) Posterior distributions of *α*_1_ and *α*_2_ comparing dynamical models for each RF.

As with the white-noise case, the estimated RFs were qualitatively similar to the input RFs, but with distortions in the temporal profile. Most of the estimated RFs appeared to be smeared in time and with stronger and longer suppressive periods. Some of the distortions were consistent across tonic and phasic models, but there were also differences between the two dynamical models that reflect the effects of K_LT_. We analyzed these effects by looking at the tMTFs, which are calculated by summing the 2D Fourier transform of the RFs across the spectral dimension (Fig. 4G). These plots show how well the model neuron is able to encode temporal modulations in the stimulus as a function of frequency. All of the estimated RFs were tuned to frequencies below 100 Hz, which is about the fastest temporal modulation rate found in zebra finch song (Singh and Theunissen, 2003). Although some of the input RFs had the potential to represent faster modulations, these frequencies were attenuated in the estimated RFs, probably because of the passive filtering properties of the membrane and the statistics of the stimulus. The main differences between the dynamical models were in the attenuation of low frequencies. Strikingly, the effects of the dynamics on lowpass attenuation varied across RFs. For the WB input, the estimated tMTF was more bandpass in the phasic model compared to the tonic model, while the opposite was true for the BP-L and BP-H inputs. Thus, not only does K_LT_ change the temporal encoding properties of the neuron, but this effect is different depending on the filtering properties of the inputs (i.e., the input tMTF).

The posterior distributions for the spike-history parameters were broader than for the white-noise examples (Fig. 4H), indicating that the estimates are more poorly constrained by the data. This was expected, given that the stimulus was shorter and more correlated. Nevertheless, there was essentially no overlap between the posterior distributions for the tonic and phasic versions of any of the example models, indicating that the GLM spike-history parameters were sensitive to the biophysical dynamics. Furthermore, as the next section will show, the trends in these examples were consistent across the larger sample of RFs.

As with the RF temporal structure, the spike-history filter parameters were affected by the interaction of RF type and dynamics. In general, phasic models had stronger short-timescale adaptation than tonic models, as indicated by larger values of *α*_1_ (Fig. 4F). This effect was in the opposite direction from what we saw in the white-noise case (Fig. 3E), where tonic neurons had larger values of *α*_1_ and *α*_2_. This discrepancy presumably reflects differences in the stimulus statistics, because the white-noise example RF was qualitatively similar to the temporal profile of the example RFs. As has been reported previously, neuron models fit to white-noise stimuli produce poor predictions to natural stimuli (Theunissen et al., 2001). The white-noise GLMs produced good predictions because they were fit and tested with white-noise stimuli, but the parameter estimates do not generalize to other kinds of stimuli. As noted above, a key feature of birdsong is that the temporal envelope is dominated by low frequencies. These slow oscillations produce sustained periods of excitation or inhibition that drive the dynamical model into regimes where adaptive processes come more strongly into play. This nonlinear interaction between stimulus statistics and dynamics likely also explains why the effect of K_LT_ varied across the example RFs: phasic dynamics (i.e., increased K_LT_) caused *α*_1_ to increase for all three RFs, but only affected *α*_2_ for the BP-L RF.

### Interaction of intrinsic dynamics and RF temporal filtering

Based on these examples, we hypothesized that the key contributor to these interactions was the temporal profile of the input RF, in particular whether there was a negative lobe at longer lags. In the modulation frequency domain, this lobe corresponds to bandpass filtering. The parametric, Gabor-based model we used to generate the RFs (Woolley et al., 2009) represents this feature by a single parameter, the temporal phase (*P_t_*), which is 0 for the WB example and 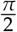 for the BP-L and BP-H examples. Approximately half (26/60) of the RFs in our larger sample, those with modulation power primarily along the spectral axis, had *P_t_* of 0, whereas the RFs with power along the temporal modulation axis (34/60) had *P_t_* of 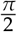.

The performance of GLMs fit to data from the larger set of RFs was consistently good, with high correlations between the spike-rate histograms of the LDC and GL models for the tonic_WB_ (*r* = 0.86 ± 0.04), tonic_BP_ (*r* = 0.90 ± .04), phasic_WB_ (*r* = 0.75 ± 0.08), and phasic_BP_ (*r* = 0.87 ± 0.05) groups, that were comparable to the correlations between the even and odd trials of the LDC data for the tonic_WB_ (*r* = 0.93 ± 0.01), tonic_BP_ (*r* = 0.84 ± 0.03), phasic_WB_ (*r* = 0.92 ± 0.02), and phasic_BP_ (*r* = 0.90 ± 0.02) models. Performance was slightly lower for the phasic_WB_ data, but the reason for this was not clear.

The results from the larger sample of RFs were consistent with our hypothesis. We looked first at the effects of dynamics on RF temporal structure, specifically the extent to which the estimated tMTF (which represents how the full LDC model encodes stimuli) was attenuated at low frequencies compared to the input tMTF (Δ*l*). In Fig. 4G, Δ*l* corresponds to the difference between the black line and blue or yellow line at *f* = 0 with maximum power set to 1. Positive values of Δ*l* indicate that the estimated RF is more bandpass (i.e., responds less to low-frequency modulations) compared to the input RF. Negative values indicate that encoding of lower frequencies is boosted. As shown in Fig. 5, for models with WB temporal tuning, phasic dynamics attenuated low frequencies, in comparison to the matching tonic models (LMM: *b*_0_ = 0.02, *b*_1_ = −0.11, *n* = 52). For neurons with BP temporal tuning, the effect was the opposite: phasic dynamics caused low frequencies to be less attenuated compared to the matching tonic models (*b*_0_ = −0.05, *b*_1_ = 0.15, *n* = 68). In other words, across a broad range of RFs, K_LT_ consistently causes neurons with broadly tuned inputs to become more selective for higher-frequency features, but causes neurons that already have narrowly tuned inputs to become more responsive to lower frequencies.

**Fig. 5.**
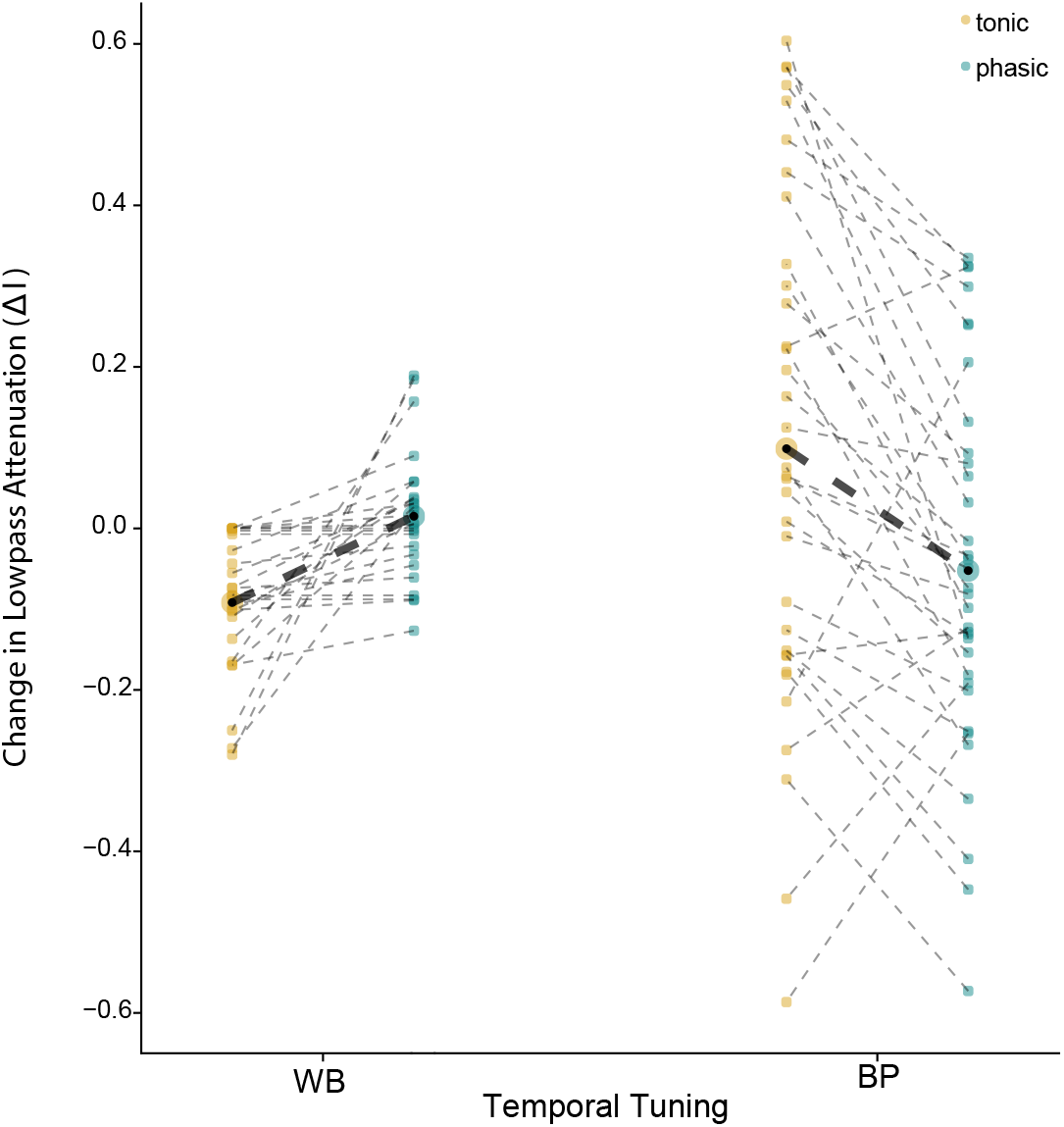
Phasic dynamics attenuate low-frequency modulations for temporal wideband RFs but enhance them for bandpass RFs. Lowpass attenuation was defined as the difference in the ratios between the power at *f* = 0 and the peak power of the temporal modulation spectrum (as in Fig. 4H) of the input RF and GLM estimated RF (Δ*l*; see Methods). The y-axis shows the difference between this value for the input RF and the estimated RF. Positive values indicate that the estimated RF is more bandpass in its temporal filtering properties compared to the input RF, while negative values indicate the estimated RFs were more lowpass. For each RF, lowpass attenuation estimates for the phasic and tonic models are connected by a black dotted line. The bold dotted line shows the differences in the mean lowpass attenuation estimates (enlarged black dot) between RF types for a given model. The linear mixed effects model (LMM) with the interaction between RF type and dynamics fits significantly better than the LMM with main effects only (LMM: *χ*^2^(1) = 19.04, *p* < 0.001)

Similarly, just as we saw with the example models, the adaptation parameters also depended on RF temporal structure and dynamics. As shown in Fig 6A, the general trend was for phasic models to have lower spontaneous firing rates and stronger adaptation, but there were some differences in the effect of phasic dynamics on *α*_2_ that depended on RF type. Models with phasic dynamics had lower baseline firing rates (larger values of *ω*; Fig 6B) compared to tonic models (LMM: *b*_0_ = 9.08, *b*_1_ = −1.29, *n* = 120), and models with WB RFs had lower baseline rates compared to models with BP RFs (*b*_2_ = −2.37, *n* = 120, Fig. 6B). Similarly, models with phasic dynamics had stronger short-term adaptation (*α*_1_; Fig. 6C) compared to tonic models (*b*_0_ = 196.77, *b*_1_ = −150.99, *n* = 120), and models with BP RFs had stronger adaptation than models with WB RFs (*b*_2_ = 1.21, *n* = 120). For both of these parameters, there was not a significant interaction between model dynamics and RF type. However, there was an interaction for longer-timescale adaptation (*α*_2_; Fig 6D). For WB RFs, *α*_2_ was larger for phasic models compared to tonic models (*b*_0_ = 0.29, *b*_1_ = −0.49, *n* = 52), but for BP RFs, *α*_2_ was larger for tonic models (*b*_0_ = −0.48, *b*_1_ = 0.19, *n* = 68). Note that in contrast to the white-noise example, *α*_2_ estimates were sometimes negative, which corresponds to a baseline facilitation (i.e., past spikes are associated with an increased probability of firing).

**Fig. 6.**
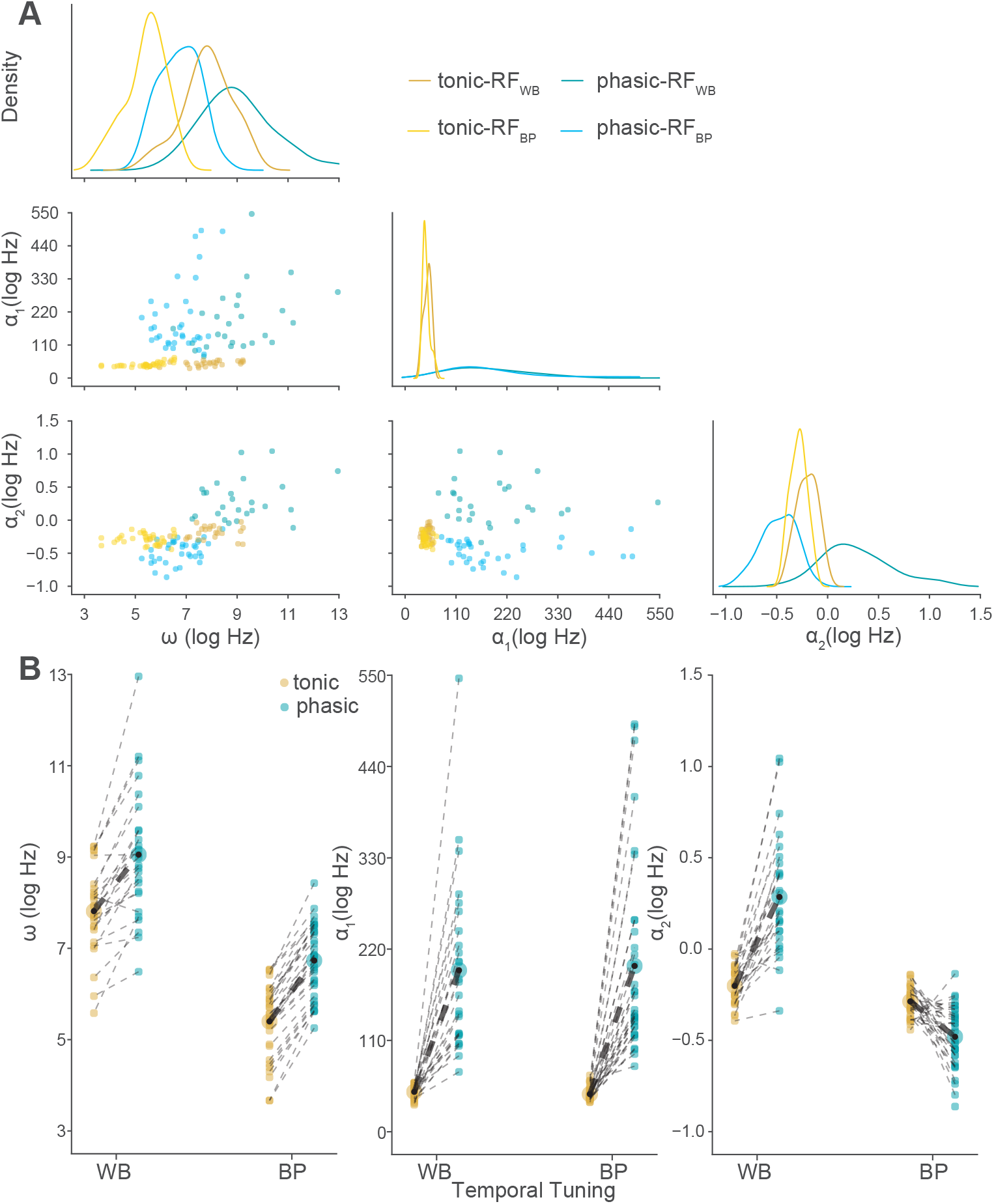
Firing rate and spike-history parameter estimates depend on RF structure and dynamics. (A) Point estimates of *ω*, *α*_1_, and *α*_2_ GLM parameters for phasic (blues) and tonic (yellows) models by RF type. Across the diagonal are the marginal distributions for each of the parameters, with the joint distributions on the off-diagonal. (B) Strip plot of parameter estimates showing paired phasic and tonic models (as in Fig. 5). For each RF, the phasic and tonic model parameter estimates are connected by a black dotted line. The bold dotted lines show the differences in the mean parameter estimates between RF types for a given model. The LMM with main effects and an interaction was a significantly better fit than an LMM with main effects only for *α*_2_ (*χ*^2^(1) = 72.00, *p* < 0.001), but not for *ω* (LMM: *χ*^2^(1) = 0.38, *p* = 0.54) or *α*_1_ (*χ*^2^(1) = 0.08, *p* = 0.78)

### Nonlinear, nonmonotonic effects of KLT on encoding properties

Up to this point, intrinsic dynamics have been dichotomized into tonic and phasic firing. For step currents, this dichotomy reflects a bifurcation in the dynamics: below a critical value of *g_KLT_*, spiking is repetitive, but above this value, it occurs only at the stimulus onset (Rothman and Manis, 2003; Meng et al., 2012). For broadband current stimuli, however, the effects of *g_KLT_* are more graded (Chen and Meliza, 2018). To test whether K_LT_ affects encoding properties in a continuous or binary manner, we simulated responses using LDC models with values of *g_KLT_* that varied in steps of 1 nS over a range of 0 to 50 nS (with capacitance kept constant at 60 pF), which encompasses the bifurcation in this model from tonic to phasic firing. For simplicity, we used only the three example receptive fields shown in Fig. 4 (WB, BP-L, and BP-H). Using the same birdsong stimulus, we fit GLMs to data from these simulations and examined how lowpass attenuation and adaptation were affected.

The correlation between even and odd trials of the simulated data tended to increase with *g_KLT_* (Fig. 7A), which is consistent with our previous finding that K_LT_ makes spike timing more precise and less variable across trials (Bjoring and Meliza, 2019). In contrast, although the performance of the GLM was good across all levels of *g_KLT_* (Fig. 7B), it tended to decrease with larger *g_KLT_* values. This suggests that the LDC model is more difficult to approximate with a GLM as additional voltage-gated conductances are added. Overall, the predicted spike trains remained highly accurate, allowing resulting parameter estimates to be meaningfully interpreted.

**Fig. 7.**
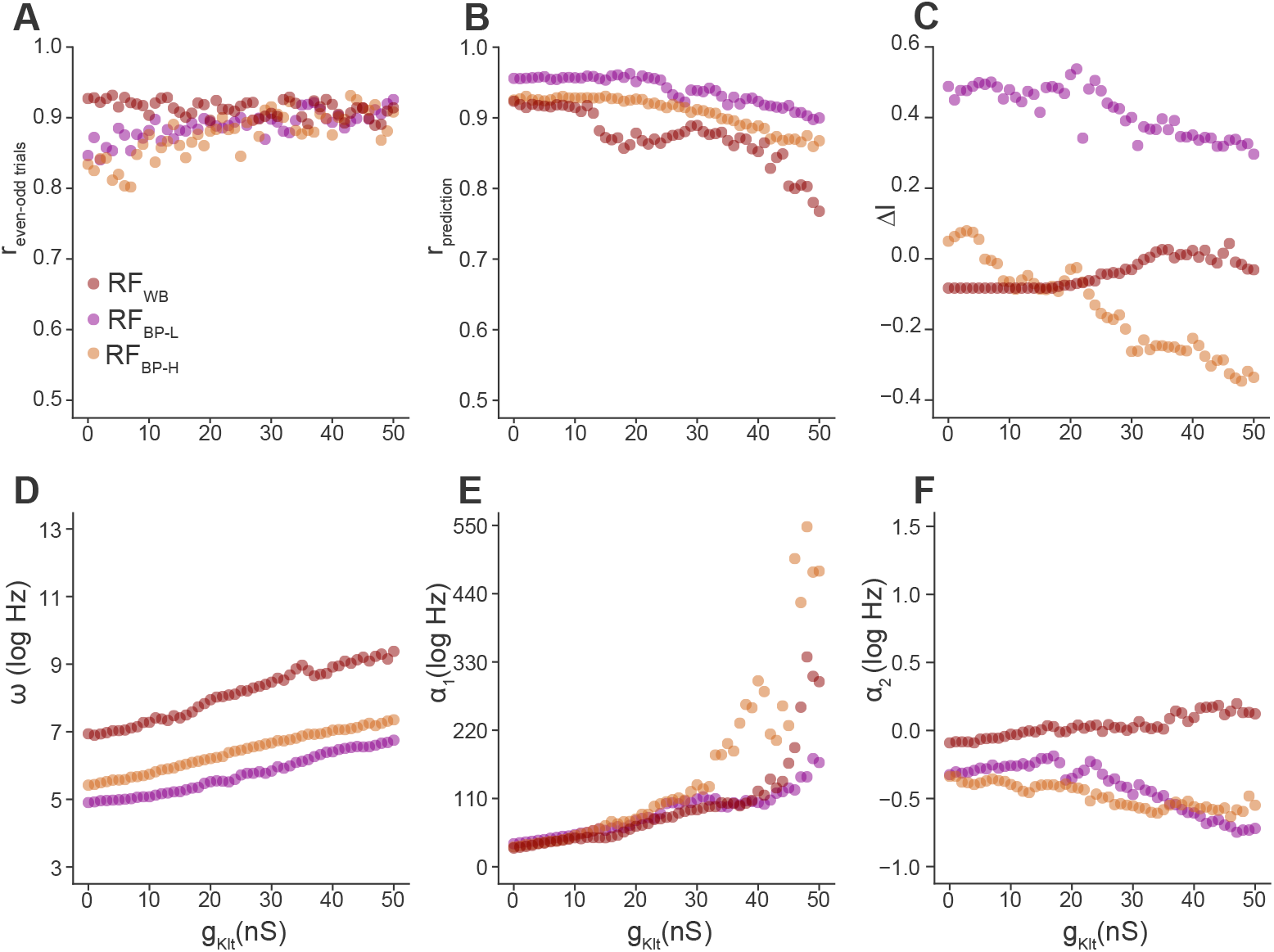
Effects of low-threshold potassium conductance (*g_KLT_*) on GLM parameters are nonlinear and depend on RF structure. (A) Correlation coefficients between the even and odd trials of the LDC model as a function of *g_KLT_* for the three exemplar RFs. (B) Correlation coefficients between the spike-rate histograms of the LDC and GL models as a function of *g_KLT_*. (C–F) Lowpass attenuation, *ω*, *α*_1_, and *α*_2_ estimates as a function of *g_KLT_*.

Consistent with what we observed with dichotomized dynamics, the effects of K_LT_ on RF temporal structure, spontaneous firing rate, and adaptation depended on RF type (Fig. 7C–F). With the exception of spontaneous firing rate (Fig. 7D), the trajectories of the parameters as *g_KLT_* increased were nonlinear although approximately monotonic. However, there was little evidence of bifurcation, which would have appeared as a sharp discontinuity between two stable regimes. These results confirm that the effects of intrinsic dynamics on encoding properties are highly nonlinear, with a strong dependence on the statistics of the stimulus and the tuning of the inputs.

## Discussion

These data demonstrate how intrinsic dynamics can affect the temporal encoding properties of cortical-level auditory neurons. Although this effect is not unexpected, to our knowledge it has not yet been quantitatively characterized. Our approach was to simulate zebra finch auditory responses with a biophysically realistic linear-dynamical cascade model and then estimate encoding properties using GLMs, which are statistically robust and easy to interpret. This allowed us to modulate intrinsic dynamics by changing the parameter values that correspond to specific cellular mechanisms and explore the effects on receptive fields and spike-history adaptation.

We focused on a low-threshold potassium current (K_LT_), which is expressed in a subset of neurons in zebra finch CM. In a previous study, we used broadband current injections to show that K_LT_ affects temporal integration, causing neurons to become more coherent with inputs at frequencies around the maximum temporal modulation rate of zebra finch song (Chen and Meliza, 2018). This effect is reproduced by the dynamical model used here. However, the current stimuli used to build the model were artificial and unrepresentative of the stimulus-driven synaptic activity CM neurons would receive in vivo. Thus, to predict how variation in K_LT_ might affect auditory responses to vocal communications in this species, we drove the dynamical model with an injected current that was the result of convolving natural zebra finch song with a spectrotemporal RF, which we term the “input RF”. Input RFs, which represent a linear approximation of the processing performed by the neuron’s presynaptic partners and the dendritic integration of excitatory and inhibitory synaptic currents, were randomly drawn from a published distribution of RFs found in zebra finch Field L (Woolley et al., 2009), the major source of ascending auditory input to CM (Vates et al., 1996; Wang et al., 2010). This allowed us to predict which effects of the dynamics would be consistent across the population and which would depend on tuning of the inputs.

### K_LT_ has a nonlinear influence on how neurons encode stimuli

The estimated RFs, which we interpret as the features of the stimulus that neurons encode in their spiking outputs, reflected the statistics of the stimulus, the filtering properties of the input RFs, and the dynamics of spiking. Estimated RFs qualitatively resembled input RFs but were distorted in time. Analyzing these distortions using temporal modulation transfer functions (Fig. 4), we found that most (71/120) of the model neurons were less responsive to high frequencies (≥ 100 Hz) than their inputs; we expected this effect from the lowpass filtering associated with passive leak currents. K_LT_, in contrast, primarily affected low frequencies in the tMTF. To our surprise, the sign of the effect depended on the input tMTF, specifically how broadly tuned it was. Wideband tMTFs became more bandpass, with stronger attenuation at low frequencies. Bandpass tMTFs, however, became more lowpass, indicating that K_LT_ was effectively boosting responses to low frequencies in the stimulus.

This result is somewhat counterintuitive, but it is consistent with the high degree of nonlinearity phasic neurons exhibit for low-frequency inputs. Using slice recordings, we previously showed that phasic and tonic CM neurons differ in their coherence between current input and spiking output (Chen and Meliza, 2018), with phasic neurons exhibiting lower coherence than tonic neurons for frequencies below about 20 Hz. Because ideal linear time-invariant systems have coherence values equal to unity for all frequencies (Marmarelis, 1988), this result indicates that phasic neurons are more nonlinear at low frequencies, but not the sign or magnitude of the nonlinearity (*contra* our interpretation in that study). In other words, for some stimuli phasic neurons may boost low frequencies while for other stimuli they may attenuate low frequencies. This is precisely the effect we observed here.

### K_LT_ has a nonlinear influence on how neurons adapt to prior activity

K_LT_ also affected the spike-history filter component of the GLM. Here the effects were more consistent across RF types, though there was a weak but significant interaction for long-term adaptation (*α*_2_), such that WB neurons became more strongly adapting with phasic dynamics and BP neurons became more facilitating (Fig. 6B). Within the joint distribution of all the spike-history parameters (*ω*, *α*_1_, and *α*_2_; Fig. 6A), there was there was a clear visual separation in the population distributions of tonic and phasic neurons, such that one could potentially infer whether a cell was tonic or phasic from the spike-history parameters alone. Thus, under some circumstances it may be possible to use extracellular recordings to characterize intrinsic dynamics.

When dynamical neuron models are stimulated with step currents, *g_KLT_* is a bifurcation parameter with a critical value that determines whether the cell can spike repetitively (tonic firing) or not (phasic firing). We found that using more realistic currents, there is little evidence of bifurcation in encoding properties, which changed smoothly as we varied *g_KLT_* (Fig. 7). These relationships nonetheless tended to be quite nonlinear, indicating that neurons can in principle achieve dramatic changes in functional response properties with only small changes in the expression or localization of a single type of channel.

### Functional implications of K_LT_ expression in the avian auditory system

Taken together, these results demonstrate that the encoding properties of auditory neurons can be highly sensitive to changes in intrinsic dynamics arising from the inclusion or exclusion of a single current. We recently showed that CM neurons express more Kv1.1 and become more phasic during the peak of the critical period for song memorization, but only in finches raised in the complex acoustic environment of a colony (Chen and Meliza, 2020). As suggested by our results here, increased expression of a low-threshold potassium channel like Kv1.1 might help neurons to filter out this kind of background noise by selectively suppressing responses to low-frequency inputs in neurons that have broad temporal tuning. Such a mechanism could explain the recent finding that in rats that exposure to dynamically modulated noise causes neurons in primary auditory cortex to shift their tuning away from the spectrotemporal modulation frequencies of the noise (Homma et al., 2020). In this respect, K_LT_ may be serving an analogous function to the co-tuned feedforward inhibitory inputs seen in mammalian auditory cortex (Wehr and Zador, 2003; Tan et al., 2004), but without the need for a separate population of neurons. As we have speculated elsewhere, a cell-intrinsic mechanism for filtering out background noise and increasing spike precision may be an important complement to synaptic plasticity early in development when inhibitory circuits and the reversal potential of inhibitory conductances are still stabilizing (Chen and Meliza, 2020).

It is less clear to us why it would be useful for K_LT_ to boost low-frequency responses in neurons that already have bandpass-tuned inputs; however, we note that this effect was considerably more variable (compare the variance for BP and WB neurons in Fig. 5). Moreover, it is not yet known if the distribution of K_LT_ expression in CM is independent of the distribution of input temporal tuning. If expression of K_LT_ depends on experience, and more proximately on the statistics of presynaptic and postsynaptic activity, then its effects may be restricted to neurons with specific tuning properties. Intracellular recordings to measure excitatory and inhibitory RFs in CM neurons may be needed to determine if this is the case.

### Model-based approaches to understanding how nonlinear mechanisms affect sensory processing

This study complements other efforts to incorporate biologically realistic mechanisms into the framework of linear-nonlinear cascade models. Early work in the auditory system demonstrated how static nonlinearities in the summation of RF components alter the encoding properties of stochastic spiking models Christianson et al. (2008). More recent studies have added idealized representations of dynamical mechanisms like excitatory and inhibitory conductances (Schinkel-Bielefeld et al., 2012) or gain adaptation Ozuysal and Baccus (2012) to the linear-nonlinear framework, while retaining the ability to statistically estimate the parameters of these model components and use them to predict biological data. In comparison, our approach emphasizes realism, building on a detailed biophysical model of intracellular voltage dynamics with pharmacologically and (in principle) genetically identifiable components. This realism comes at the cost of statistical tractability. We have addressed this issue by using an entirely different but much simpler model to characterize the encoding properties of the more complex model. Although this limits us to asking empirical questions, there are many biological insights to be gained from an empirical approach.

Within this biophysically realistic framework, our analysis was limited to the effects of manipulating a single biophysical parameter (*g_KLT_*) on encoding of a single kind of auditory stimulus (zebra finch song). It is important to note that the nonlinearity of neuronal dynamics means that our results are therefore only valid within the specific context of the other ionic currents in the model. In a different cell type that expresses a different complement of currents, K_LT_ will interact with those currents differently and may have entirely different effects on sensory coding. However, although the results may not generalize broadly, the approach can be adapted widely, to other auditory areas and sensory systems that exhibit diverse or plastic intrinsic dynamics. We have shown that GLMs can accurately predict the spiking responses of more complex, more biophysically realistic models across different kinds of stimuli, receptive fields, and dynamical regimes. Care is needed in interpreting the GLM parameter estimates, which do not correspond to specific cellular mechanisms and are therefore not linear or independent functions of the underlying dynamics. Given the nonlinear kinetics of most voltage-gated currents, we expect that the relationships between intrinsic dynamics and encoding properties will be complex and often counterintuitive in most systems, but that there will be much to learn in each system about how intrinsic dynamics reflect the computational tasks and constraints that need to be solved.

## Methods

### Stimulus Design

For univariate white-noise models, the stimulus consisted of 100 s of Gaussian white noise sampled at 1 kHz. For multivariate models, the stimulus consisted of zebra finch song motifs recorded from 30 adult males in our colony. Each motif was normalized to the same RMS amplitude and repeated twice, padding with at least 50 ms microphone noise at the beginning to avoid transients in the convolution. The total duration of the stimulus was 63.7 s, of which 12.7 s was reserved for testing performance. Spectrograms of the stimuli were calculated using a gammatone filter bank (Slaney, 1998) with a window size of 2.5 ms and 20 spectral channels between 1.0 and and 8.0 kHz, and a step size of 1.0 ms.

### Receptive Field Construction

The univariate white-noise receptive field was generated from the difference of two gamma functions 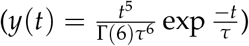 with time constants of 16 and 32 ms and an amplitude ratio of 1.5. Spectro-temporal receptive fields (RFs) were parameterized as the outer product of two Gabor functions multiplied by a scalar amplitude:

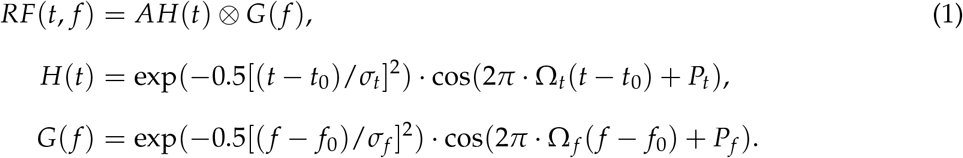

where *H* is the temporal dimension of the RF, *G* is the spectral dimension, *t*_0_ is the latency, *f*_0_ is the peak frequency, *σ_t_* and *σ_f_* are the temporal and spectral bandwidths, Ω_*t*_ and Ω_*f*_ are the temporal and spectral modulation frequencies, *P_t_* is the temporal phase (either 0 or 2*π*), *P_f_* is the frequency phase (set to 0 for all RFs), and *A* is the amplitude. The temporal dimension *H* had a duration of 50 ms with a 1 ms resolution, while the frequency dimension *G* had 20 channels between 1 and 8 kHz. We generated 60 RFs by sampling randomly from the distributions given in (Woolley et al., 2009) as representative of empirically recorded RFs in primary areas of the zebra finch auditory pallium. The amplitude parameter *A* was initially set to 1 for all of the RFs, but was adjusted to between 1.5–6 for 8/60 models so that they would fire at least at 1 Hz on average.

### Linear-Dynamical Cascade Model

Auditory responses were simulated with a model consisting of a linear, time-invariant stage whose output serves as an external driving current *I*(*t*) for a conductance-based, single-compartment dynamical stage (Bjoring and Meliza, 2019).

The linear stage consists of a time-invariant receptive field (RF) that is convolved with the stimulus. For the univariate white-noise stimuli, this was a simple 1-dimensional convolution. For the song stimuli, each spectral channel was convolved with the corresponding channel of the RF and the results were summed to produce a univariate time series. In each trial, the output of the convolution *D*_stim_(*t*) was added to a randomly generated signal *D*_noise_(*t*) with a spectral power distribution of 1/ *f* and a signal-to-noise ratio of 4. The total drive *D*(*t*) = *D*_stim_(*t*) + *D*_noise_(*t*) was unbounded. For the white-noise stimuli, this was not an issue, and drive was converted to current *I*(*t*) with a constant scaling factor. However, for song stimuli *D*(*t*) often reached unrealistic values. Because spectral power is always positive, RFs with lowpass temporal characteristics tended to over-drive the neurons with long periods of net positive current. Given that excitation and inhibition are generally balanced in the mammalian auditory cortex (Wehr and Zador, 2003), and that synaptic currents in biological neurons are limited by the reversal potentials of sodium, potassium, and chloride, for song stimuli we therefore mean-centered *D*(*t*) and compressed the resulting drive to obtain a more realistic current *I*(*t*):

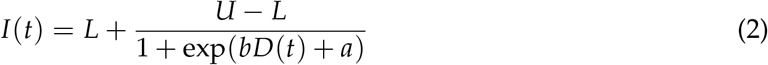

where *U* and *L* are the upper and lower bounds of input current respectively and free parameters *b* and *a* control the slope and intercept of the logistic curve. *U* and *L* were calculated based on the passive membrane properties of the model such that the model would not be driven above 0 mV or below −100 mV, resulting in *U* = 97.5 pA and *L* = −32.5 pA. The free parameters were estimated by minimizing the mean squared error between Eq 2 and the identity function rectified at *U* and *L* to give *b* = −0.04 and *a* = 1.32. We also ran all the analyses without mean-centering and compression. The results were qualitatively similar, indicating that the model is robust to assumptions about the strength of the driving current. However, we only report the results from the simulations with mean-centering and compression due to their increased biological realism.

The voltage dynamics were based on a model of dorsal cochlear neurons (Rothman and Manis, 2003) adapted for tonic and phasic CM neurons by Chen and Meliza (2020). The component currents include an external driving current *I*(*t*) and six intrinsic currents.

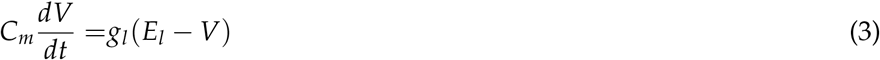

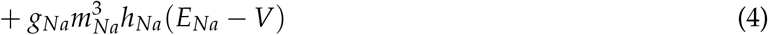

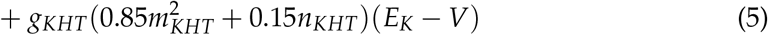

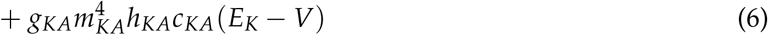

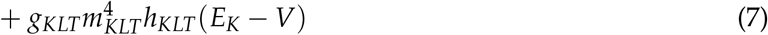

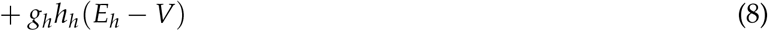

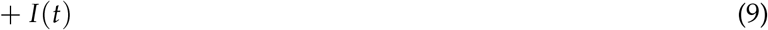

Each voltage-gated current depended on a maximal conductance *g_X_*, the reversal potential for the ion species conducted by the channel *E_X_*, and one or more gating variables (e.g., *m_X_*, *h_X_*). For all currents, the dynamics of the gating variables were defined by first-order kinetics; for example,

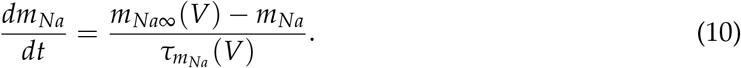

This model can produce phasic or tonic responses to step currents depending on the value of *g_KLT_*. When *g_KLT_* is low, the model neuron produces sustained responses to weak and moderate depolarizations; when *g_KLT_* is high, the model only fires at the onset of the current step. The principal model parameter values used here are shown in Table 1 (see Bjoring and Meliza 2019 for a complete list). Each RF was paired with a tonic and a phasic model. To examine how encoding properties change over the full range of *g_KLT_* values, we started with the tonic model parameters and increased *g_KLT_* from 0 nS to 50 nS in steps of 1 nS.

**Table 1.** Parameter values for biophysical models.

| Parameter | Tonic | Phasic |
| --- | --- | --- |
| $C_m$ (pF) | 60 | 40 |
| $E_l$ (mV) | -75 | -75 |
| $g_l$ (nS) | 1.3 | 1.3 |
| $E_{Na}$ (mV) | 55 | 55 |
| $g_{Na}$ (nS) | 750 | 750 |
| $E_K$ (mV) | -82 | -82 |
| $g_{KDR}$ (nS) | 0 | 0 |
| $g_{KHT}$ (nS) | 95 | 95 |
| **$g_{KLT}$ (nS)** | 0 | 50 |
| $g_{KM}$ (nS) | 0 | 0 |
| $g_{KA}$ (nS) | 30 | 30 |
| $E_h$ (mV) | -43 | -43 |
| $g_h$ (nS) | 0.5 | 0.5 |

Symbols: $C_m$ , capacitance; $E_l$ , leak current reversal potential; $g_l$ , leak conductance; $E_{Na}$ , sodium reversal potential; $g_{Na}$ (maximum) sodium conductance; $E_K$ , potassium reversal potential; $g_{KDR}$ , delayed-rectifier potassium conductance; $g_{KHT}$ , high-threshold potassium conductance; $g_{KLT}$ , low-threshold potassium conductance; $g_{KM}$ , M-type (slowly activating) potassium conductance; $g_{KA}$ , A-type (slowly inactivating) potassium conductance; $E_h$ , reversal potential for h-type (hyperpolarization-activated, cation-nonselective) current; $g_h$ , h-type conductance. Bold highlights parameter differences in tonic and phasic models.

**Table 2.** Results of Δ*l* LMM comparison.

| Model | AIC | BIC | $\chi^2(\text{df})$ | p-value |
| --- | --- | --- | --- | --- |
| Variance Components | -22.33 | -13.99 |  |  |
| + Main Effects | -21.49 | -7.55 | 3.15 (2) | 0.21 |
| + Interaction | -38.53 | -21.80 | 19.04 (1) | <0.001 |

Reductions in AIC and BIC, as well as statistically significant chi-squared tests indicated that the LMM with main effects and interaction was the best candidate model for describing the data.

**Table 3.** Results of *ω* LMM comparison.

| Model | AIC | BIC | $\chi^2(\text{df})$ | p-value |
| --- | --- | --- | --- | --- |
| Variance Components | 436.95 | 445.31 |  |  |
| + Main Effects | 277.91 | 291.85 | 163.04 (2) | <0.001 |
| + Interaction | 279.53 | 296.25 | 0.38 (1) | 0.54 |

There were reductions in AIC and BIC, as well as statistically significant chi-squared tests for the main effects model compared to the variance components model. However, there were increases in AIC and BIC, as well as a non-statistically significant chi-squared test for the main effects and interaction model. Therefore, the LMM with main effects only was the best candidate model for describing the data.

**Table 4.**
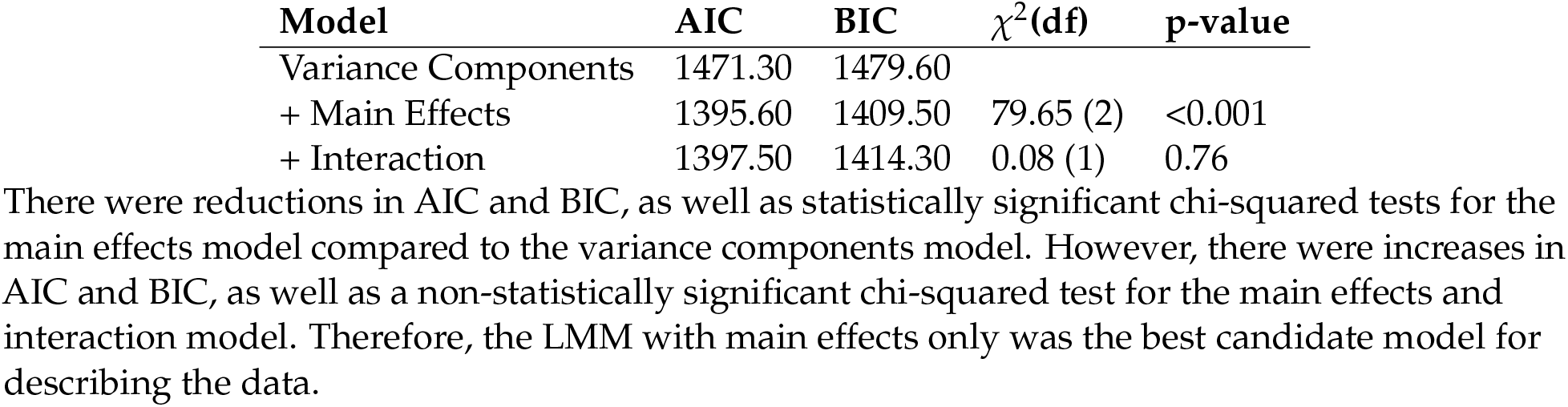
Results of *α*_1_ LMM comparison.

**Table 5.**
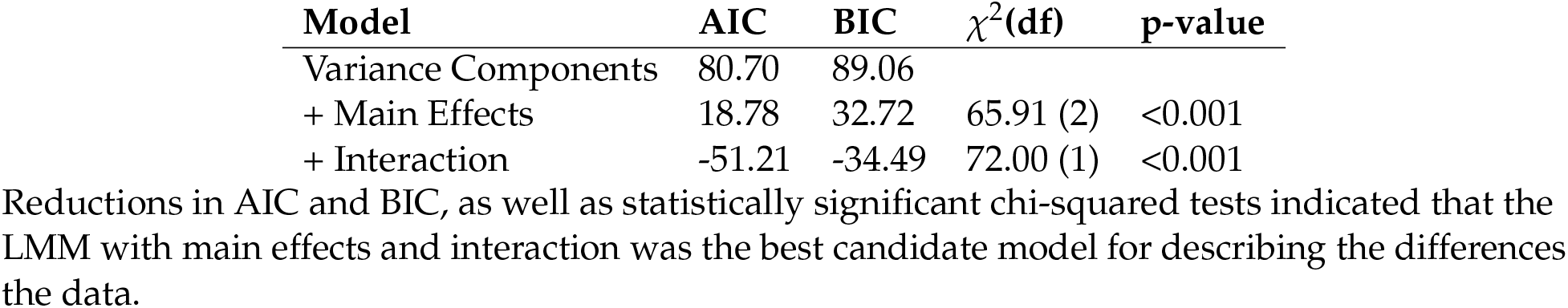
Results of *α*_2_ LMM comparison.

| Model | AIC | BIC | $\chi^2(\text{df})$ | p-value |
| --- | --- | --- | --- | --- |
| Variance Components | 80.70 | 89.06 |  |  |
| + Main Effects | 18.78 | 32.72 | 65.91 (2) | <0.001 |
| + Interaction | -51.21 | -34.49 | 72.00 (1) | <0.001 |

Reductions in AIC and BIC, as well as statistically significant chi-squared tests indicated that the LMM with main effects and interaction was the best candidate model for describing the differences the data.

The dynamical model simulation code was generated using spyks (https://github.com/melizalab/spyks; version 0.6.10), and the dynamics were integrated using a 5th-order Runge-Kutta algorithm with an adaptive error tolerance of 1 × 10^−5^ and an interpolated step size of 0.025 ms. The output of the integration was converted to spike times by thresholding the voltage at −20 mV.

### Generalized Linear Models

A generalized linear model (GLM) (Pillow et al., 2005; Calabrese et al., 2011) was fit to the spike trains produced by the linear dynamical cascade models (Fig. 2). The conditional intensity of the model was given by:

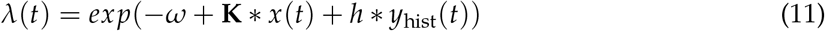

where *λ*(*t*) is the conditional intensity at time *t*, exp(−*ω*) corresponds to the baseline firing rate of the GLM, **K** is the RF, which is convolved with the song spectrogram *x*, and *h* is the spike adaptation filter, which is convolved with the spike train history *y*_hist_(*t*). Note that we use *f*_1_ ∗ *f*_2_(*t*) to denote the convolution of two functions with respect to time. The full RF was a 20 × 50 matrix (20 spectral channels by 50 time bins of 1 ms). To reduce the number of parameters and avoid overfitting, **K** was parameterized with a rank-2 approximation; that is, the product of a 20 × 2 spectral filter and a 2 × 50 temporal filter (Thorson et al., 2015). The parameter count in the temporal dimension was further reduced by projecting into a basis set consisting of 12 raised cosine functions (Pillow et al., 2005). This basis set achieves good temporal resolution in the time immediately following a spike, with the resolution smoothly decreasing at long time intervals. The spike-history filter *h* was parameterized in a basis set of two exponential functions:

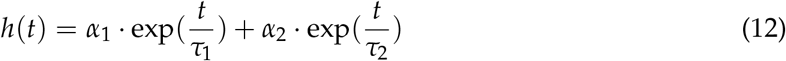

where *τ*_1_ and *τ*_2_ are time constants corresponding to short (10 ms) and long (200 ms) timescales, and *α*_1_ and *α*_2_ are the coefficients. This parameterization was chosen based on the multiadaptive timescale model, which is closely related to the GLM and has been shown to be capable of reproducing a broad range of intrinsic dynamics (Kobayashi et al., 2009; Yamauchi et al., 2011) The GLMs were fit to data from the first 80% of the stimuli. The log-likelihood function of the GLM is given by

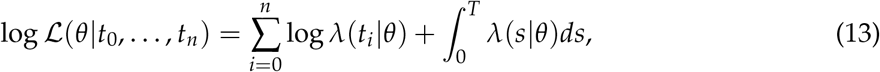

where *t_i_* is the time of the *i*th spike, *n* is the number of spikes in the experiment, *T* is the final time point of the experiment, and *θ* represents the free parameters (Rasmussen, 2018). Because the stimulus is highly correlated and the RF is expected to be sparse, we used elastic-net regularization to constrain the RF parameter estimates. Elastic-net regularization is combination of ridge regression and the least absolute shrinkage and selection operator (LASSO). Ridge regression introduces an *L*_2_ penalization parameter (*ν*_2_) to account for multicollinearity, which is inherently present in the highly correlated structure of the song spectrogram. The LASSO introduces an *L*_1_ penalization parameter (*ν*_1_) to shrink small correlations to zero and acts as a feature selection algorithm, enforcing RF sparseness. A cost function was given by:

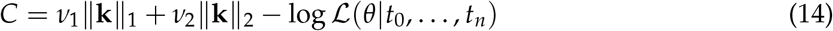

where ||**k**||_1_ and ||**k**||_2_ are the L1-norm and L2-norm of **K** (reshaped into a 1-D vector), respectively. Since the log-likelihood function is concave and is guaranteed to be free of local maxima (Paninski et al., 2004), we simultaneously estimated the parameters (*ω*, **K**, *α*_1_, *α*_2_) by minimizing the cost function, which was done by using the nonlinear conjugate gradient method scipy function ‘fmin_ncg’ (version 1.3.0) (Virtanen et al., 2020). Theano (version 1.0.4) (Al-Rfou et al., 2016) was used to symbolically derive the gradient and Hessian of the cost function and dynamically generate C code to evaluate them. The regularization coefficients (*ν*_1_, *ν*_2_) and the factorization rank *D* were chosen using 4-fold cross-validation on the estimation data.

We quantified the uncertainty in the maximum-likelihood estimates (*ω*, **K**, *α*_1_, *α*_2_) by sampling from the joint posterior distribution using emcee (version 2.2.1), a Python implementation of an affine-invariant ensemble Markov chain Monte Carlo sampler (Foreman-Mackey et al., 2013). to estimate the posterior uncertainty in the parameter estimates, *p*(*θ*|*t*_0_, …, *t_n_*) ∝ *p*(*θ*)*L*(*θ*|*t*_0_, …, *t_n_*). The log of the prior probability *p*(*θ*) was set to the elastic-net penalty (Eq. 14) using the values of *ν*_1_ and *ν*_2_ obtained through cross-validation, and the log-likelihood was as in Eq. (13). An ensemble of 1000 chains was initialized with random values centered around the maximum-likelihood estimate and given a burn-in of 2500–6000 steps. After this period, each chain was sampled one more time to give a set of 1000 independent samples from *p*(*θ*|*t*_0_, …, *t_n_*). For population-level analyses, the final value of the GLM (*ω*, **K**, *α*_1_, *α*_2_) parameters were the median value of their respective posterior distributions due to the symmetric bell-shaped curve of the posteriors. These values were very close to the initial ML point estimates, so we did not sample from the posterior for the analyses shown in Fig. 7.

To quantify performance, we generated posterior predictive distributions of spike trains from the fitted GLMs, with time discretized to Δ = 0.5 ms. At such short time scales, the conditional rate *λ*(*t*) · Δ could be approximated as a Bernoulli trial at each time bin which was used to produce spike train responses from the GLMs. In each trial, we drew a sample from the posterior distribution, so the intertrial variability reflects not only the intrinsic variance of the Bernoulli distribution but the uncertainty in the parameter estimates as well. Performance was quantified as the product-moment correlation between the spike-rate histograms (50 trials, 10 ms bins) for the data and the prediction on the 20% of the stimulus reserved for testing. As a baseline measure of intrinsic variability, we calculated the product-moment correlation between even and odd trials in the data (i.e., from the linear-dynamical cascade model); however, we did not explicitly correct performance scores.

### Lowpass Attenuation

The estimated RF parameters were projected back into a linear time basis and reshaped into a 20 × 50 matrix. To obtain the temporal modulation transfer function (tMTF), a 2-dimensional Fourier transform was performed on the RF, summing across the spectral dimension (including positive and negative frequencies). The Fourier transform was calculated using the numpy package in Python, with zero-padding and the application of a Hanning window in the temporal profile to avoid edge effects. RF lowpass attenuation was quantified as:

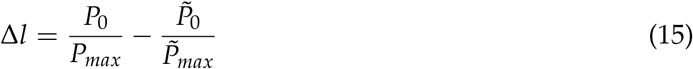

where *P*_0_ is the power for the zero frequency of the input tMTF, *P_max_* is the maximum power of the input tMTF, 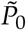 is the power at the zero frequency of the estimated tMTF, and 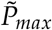 is the maximum power of the estimated tMTF. Positive values of Δ*l* indicate that the estimated RF responds more weakly to low modulation frequencies compared to the input RF, whereas negative values indicate that the estimated RF is more responsive to low frequencies.

### Linear Mixed-Effects Models

Given the nested, repeated-measures nature of the experimental design (each input RF was used with tonic and phasic dynamical models), we used a random-intercepts LMM with input RF as a random effect. All LMMs were estimated using the lme4 (version 1.1.21) R package, which does not return p-values for parameter estimates due to unreliability issues (Bates et al., 2015). To determine statistical significance, we therefore took a model-comparison approach where nested LMMs of increasing complexity were compared against each other. Three candidate models were fit: random effects (variance components) only, random effects and main fixed effects, and random effects with main effects and interactions. Restricted maximum likelihood (REML) parameter estimation gives unbiased LMM estimates, however the LMMs cannot be compared as nested models (Pinheiro and Bates, 2006) and we therefore used maximum likelihood estimation (MLE) to generate LMMs. Candidate LMMs were compared across three fit statistics: AIC, BIC, and chi-squared. Lower values of AIC and BIC indicate better relative fit. The null hypothesis of the chi-squared test is that the more complicated model is not a better fit to the data than the less complicated model.

The variance-components model was given by the equation:

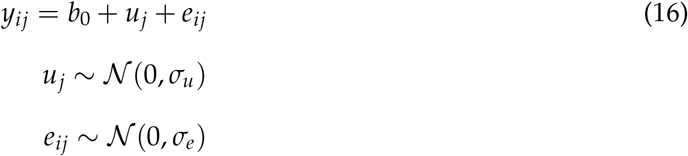

where *y_ij_* is the observed value of the dependent variable for the *i*th type of neuron model (tonic or phasic) and *j*th input RF type (WB or BP), *b*_0_ is a fixed intercept, *u_j_* is the value of the random intercept of the *j*th RF type, and *e_ij_* is the error term for the for the LMM. Both *u_j_* and *e_ij_* are assumed to be normally distributed with a mean of zero and a constant variance of 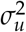 and 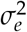 respectively. This LMM essentially tests if the differences we see in the dependent variable are solely due to the random effects of each input RF rather than neuron model or RF type. For all LMM analyses, tonic neuron models and BP RFs were coded as 1, and phasic neuron models and WB RFs were coded as 0.

The main-effects model was given by the equation:

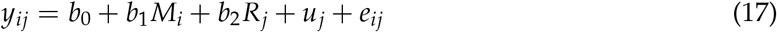

where *y_ij_*, *b*_0_, *u_j_*, and *e_ij_* are defined identically as above, *b*_1_ is the fixed effect of *M_i_*, the *i*th neuron model type, and *b*_2_ is the fixed effect for *R_j_*, the *j*th input RF type.

The interactions model is identical to the main-effects model, with the addition of a fixed effect *b*_3_ of the multiplicative interaction between neuron model and RF type, with the equation given by:

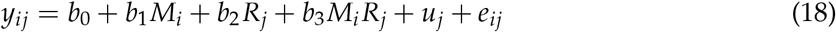

If the interactions model was found to be the best fit to the data, simple-effects models were estimated using REML since these LMMs were not compared to any other candidate models. Simple effects models were calculated by subsetting the data by RF type and estimating a LMM with RF as a random intercept and neuron model type as a fixed effect. For each RF type, the LMM equation is given by:

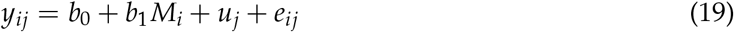

## Acknowledgments

We thank Margot Bjoring for assistance in model development and critical feedback, Laura Jamison for suggestions in statistical analysis design, and Jacy Zanussi for thoughtful discussion and critical feedback. This work was supported in part by funding from the National Science Foundation (IOS-1942480), the Thomas F. and Kate Miller Jeffress Memorial Trust, and The Hartwell Foundation.

## Supporting information captions

**S1 Text. Details of GLM estimates for exemplar tonic and phasic models shown in Fig. 4A–F.** The six figures have are in the same order as the examples in Fig. 4 and have the same format as each other: (A) Left, input RF in the LDC model. Right, estimated RF from GLM. A novel birdsong stimulus was used to compare GLM performance to output of LDC model, with 5s of the spectrogram shown in (B). Voltage traces of LDC model in response to stimulus for a single trial are shown in (C) with corresponding *K_LT_* current (D). Spike trains for all 50 trials are shown in (E), with black corresponding to the LDC model and red to the GLM. PSTHs shown in (F).

