## Supplementary material for "Nonlinear effects of intrinsic dynamics on temporal encoding in a model of avian auditory cortex": S1 Text

**S1 Text. Details of GLM estimates for exemplar tonic and phasic models shown in Fig. 4A–F.** The six figures have are in the same order as the examples in Fig. 4 and have the same format as each other: (A) Left, input RF in the LDC model. Right, estimated RF from GLM. A novel birdsong stimulus was used to compare GLM performance to output of LDC model, with 5s of the spectrogram shown in (B). Voltage traces of LDC model in response to stimulus for a single trial are shown in (C) with corresponding $K_{LT}$ current (D). Spike trains for all 50 trials are shown in (E), with black corresponding to the LDC model and red to the GLM. PSTHs shown in (F).

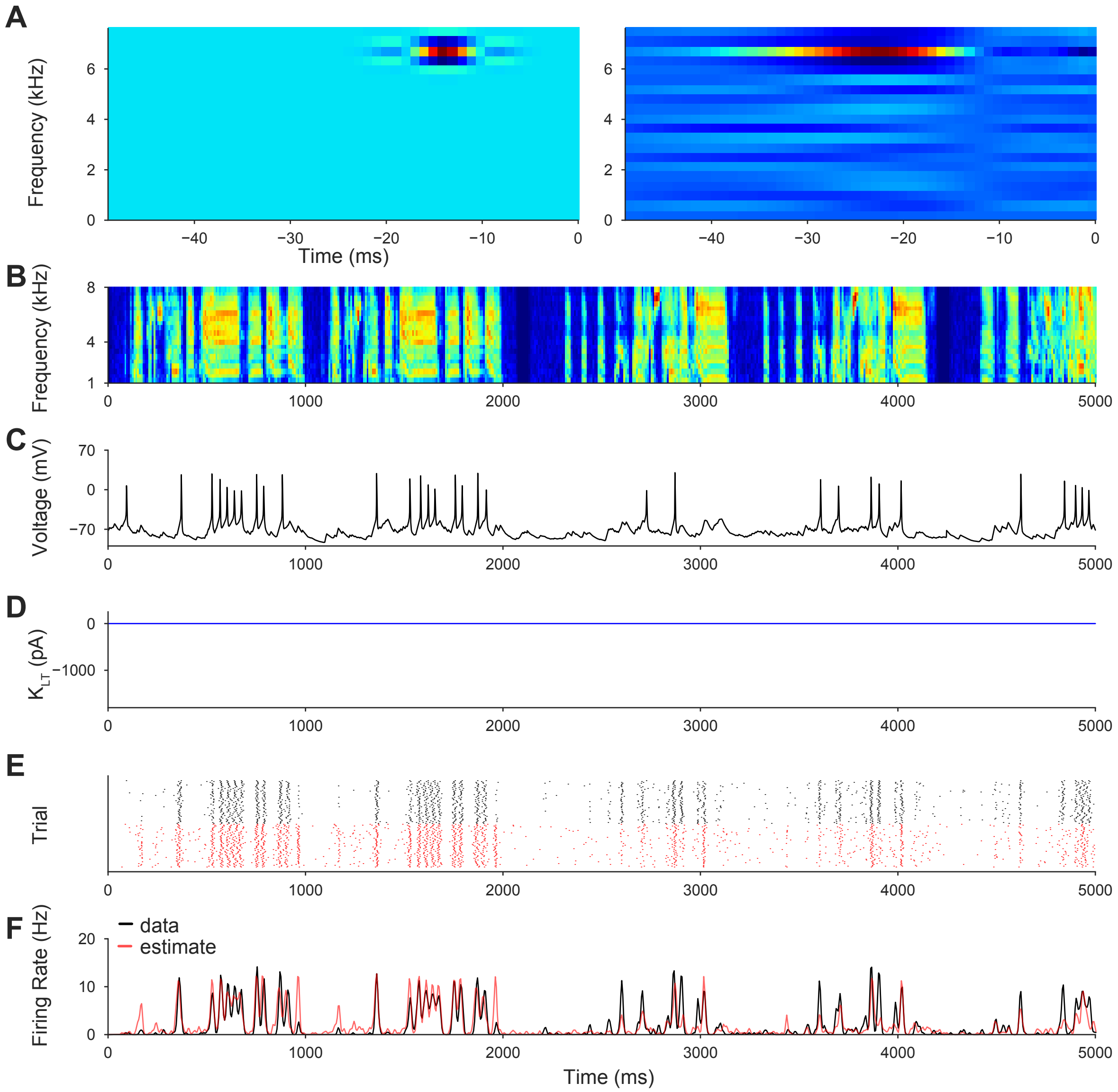


**Fig. 1. GLM estimate for tonic WB example.**

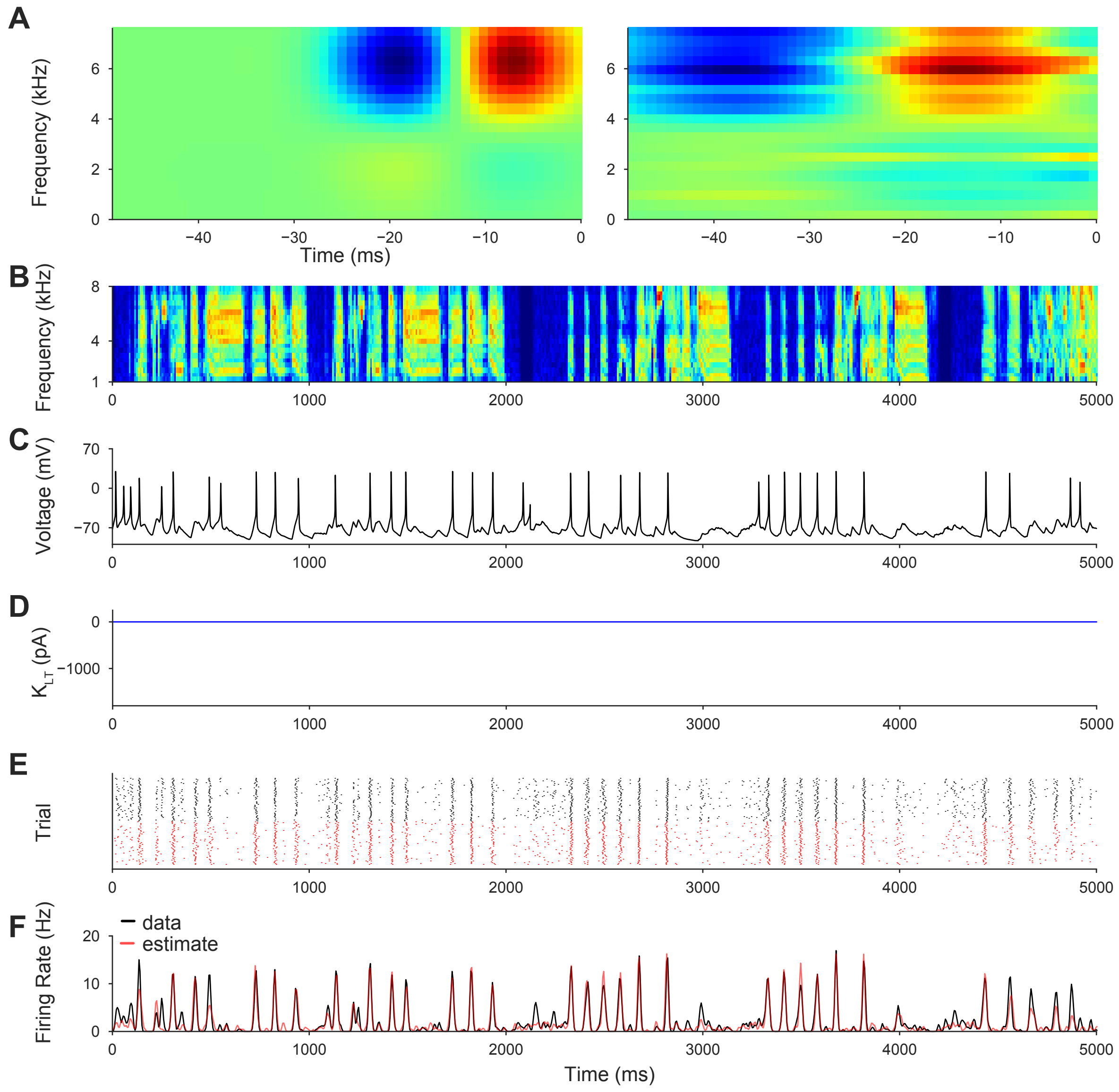


**Fig. 2. GLM estimate for tonic BP-L example.**

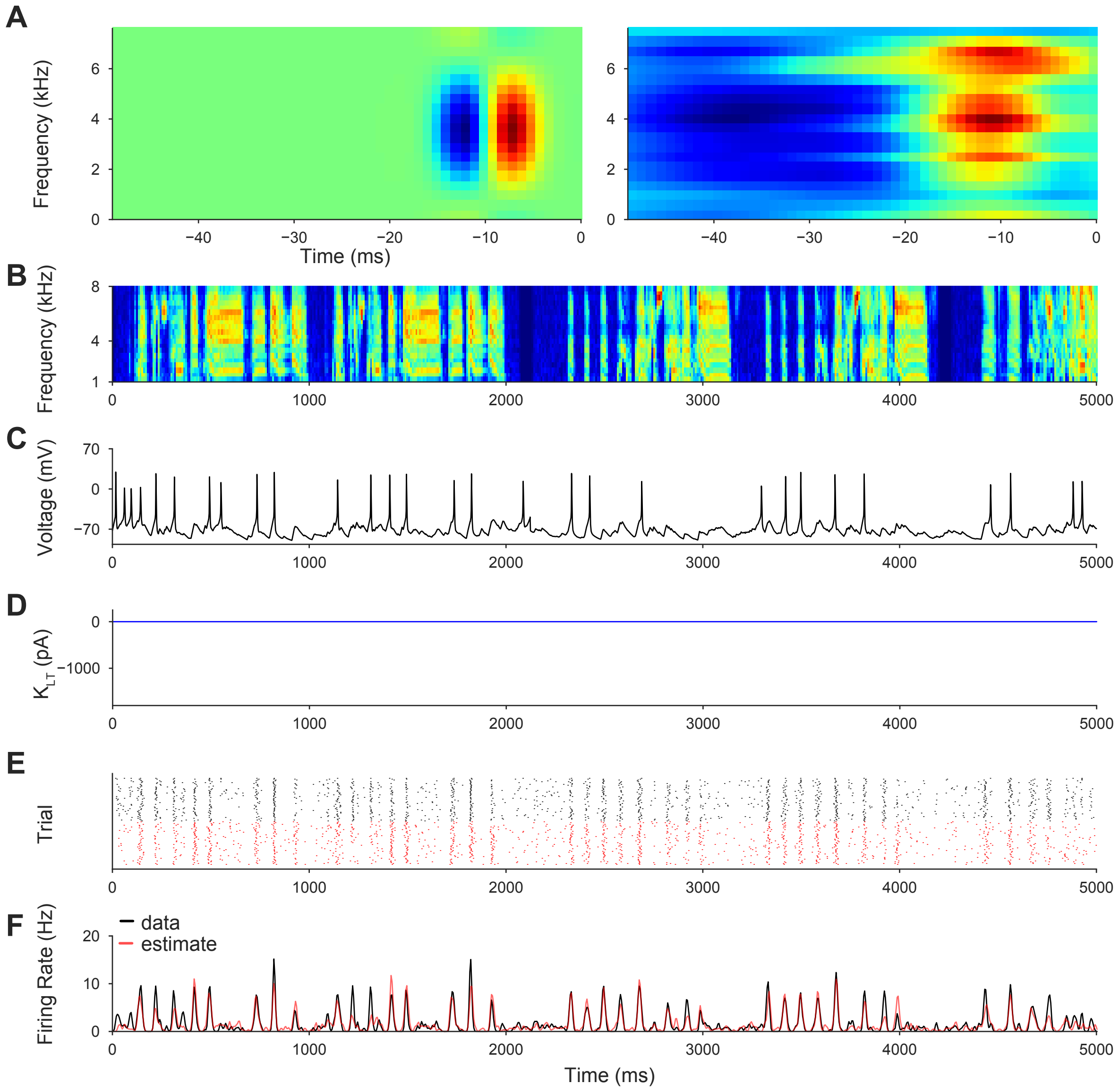


**Fig. 3. GLM estimate for tonic BP-H example.**

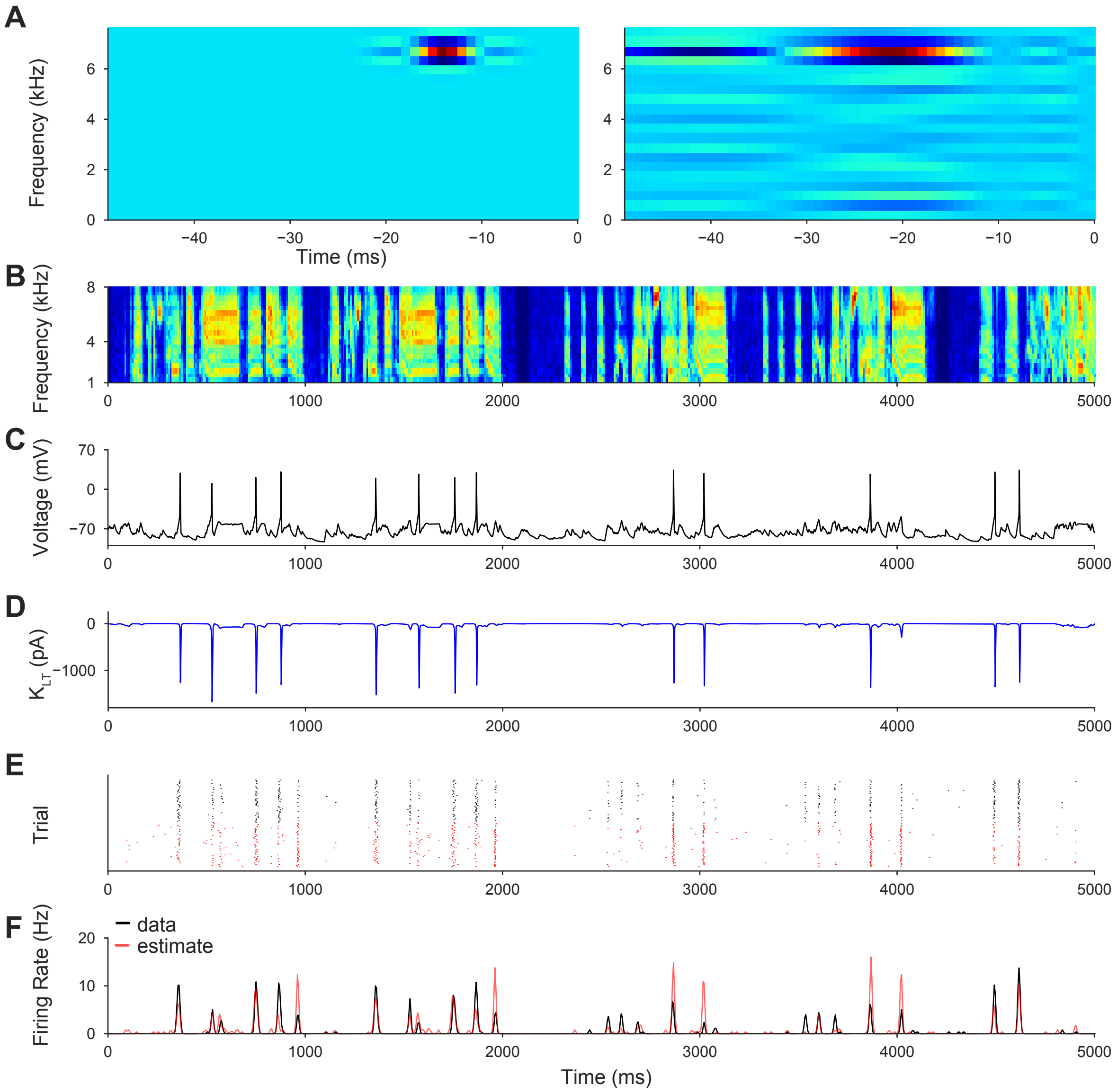


**Fig. 4. GLM estimate for phasic WB example.**

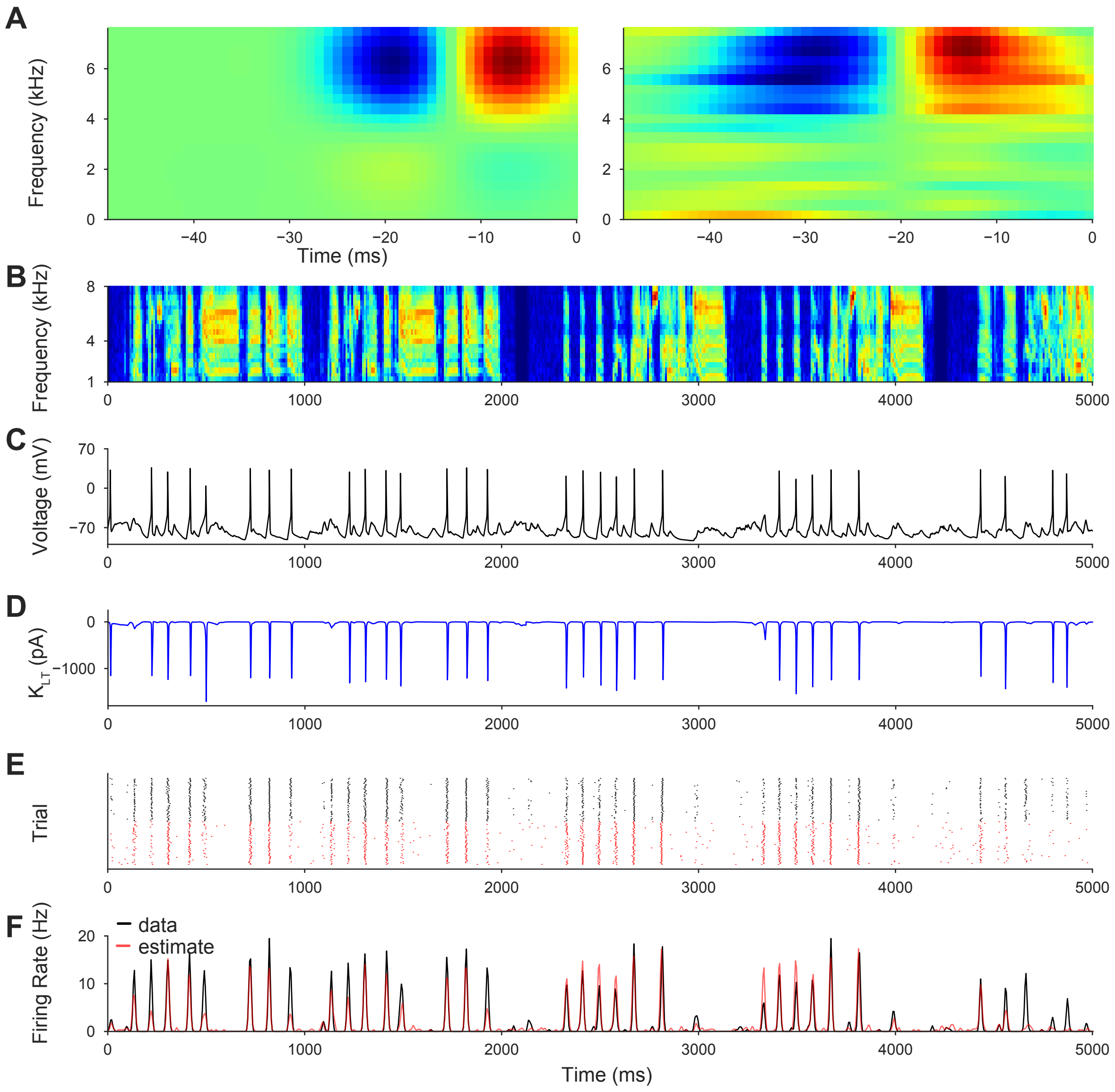


**Fig. 5. GLM estimate for phasic BP-L example.**

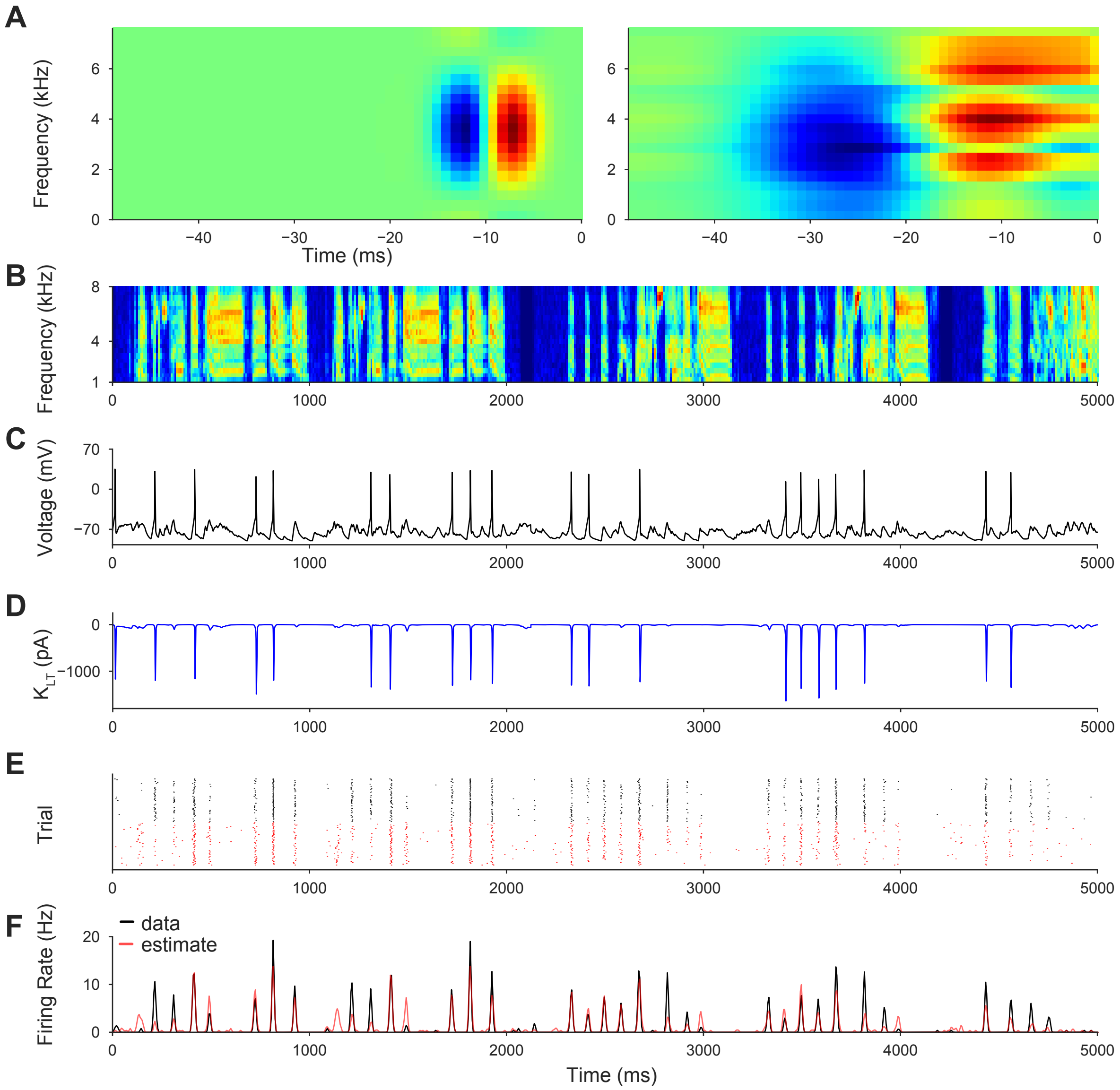


**Fig. 6. GLM estimate for phasic BP-H example.**
